# Sortilin exhibits tumor suppressor-like activity by limiting EGFR transducing function

**DOI:** 10.1101/2021.05.12.443742

**Authors:** E. Lapeyronnie, C. Granet, J. Tricard, F. Gallet, M. Yassine, A. Chermat, MO Jauberteau, F. Bertin, B. Melloni, F. Vincent, T. Naves, F. Lalloué

**Affiliations:** EA3842 CAPTuR, Contrôle de l’Activation cellulaire, Progression Tumorale et Résistance thérapeutique and Chaire de Pneumologie Expérimentale, Université de Limoges, Faculté de Médecine, 2 Rue du Dr. Raymond Marcland, 87025 Limoges CEDEX-France; Service de Pathologie Respiratoire, Centre Hospitalier et Universitaire de Limoges, 87042 Limoges CEDEX-France; Service de Chirurgie Thoracique et Cardio-vasculaire, Centre Hospitalier et Universitaire de Limoges, 87042 Limoges CEDEX-France

**Keywords:** EGFR, sortilin, *MYC*, TKI, lung adenocarcinoma

## Abstract

Lung cancer is the leading cause of cancer deaths worldwide and remains one of the most incurable. Tyrosine kinase receptors, such as the epidermal growth factor receptor (EGFR), are often aberrantly activated and drive tumor growth. Monotherapy with tyrosine kinase inhibitors to deactivate EGFR has shown initial efficacy, but their benefits tend to decline over time. EGFR acts as a transcriptional factor promoting the expression of co-oncogenic drivers, which, in turn, interact with canonical EGFR mutations to induce therapeutic relapse. This study reports that sortilin, a crucial regulator of cytoplasmic EGFR, attenuates its transducing function. Genome-wide chromatin binding revealed that sortilin interacts with gene regulatory elements occupied by EGFR. These results suggest a model, in which sortilin exhibits potential tumor suppressor-like activity by concurrently binding to regulatory elements of *cMYC*. Sortilin expression in lung adenocarcinoma may be predictive of the efficacy of anti-EGFR strategies.

---

Lung adenocarcinoma (LUAD), which is present in about ~80% of patients with non-small cell lung cancer (NSCLC), remains the leading cause of cancer deaths worldwide^1^. About 15% of these tumors contain somatic mutations in the gene encoding epidermal growth factor receptor (EGFR), constitutively activating the tyrosine kinase (TK) domain of EGFR, even in the absence of ligand stimulation. This sustained proliferative signaling^2^ creates cells in which EGFR mutants act as principal oncogenic drivers^3^. Clinically, tyrosine kinase inhibitors (TKI)^4^ limit the intensity and duration of EGFR proliferative signaling, thereby decreasing tumor aggressiveness and the course of disease^5^. However, although early and advanced LUAD do not differ in EGFR mutation frequency or type^6^, the clinical benefits of TKIs decline over time^5,7^.

Irrespective of disease stage, co-oncogenic drivers cooperate with canonical EGFR mutations in maintaining tumor malignancy and enhancing relapse. EGFR can act as a transcriptional factor^8–10^, directly promoting the expression of these co-drivers, such as *MYC* and *CCND1*, which have been implicated in epigenic reprogramming^11^ and cell proliferation, respectively. These findings suggest that exclusive of its TK activity, EGFR function may be reoriented to its nuclear signaling network. Thus, controlling the spatiotemporal distribution of EGFR remains crucial in limiting its oncogenic driving force. We have reported that sortilin, a sorting receptor belonging to the vacuolar protein sorting 10 (VSP10) family, acts as a crucial regulator of EGFR endocytosis, limiting its proliferative signaling. To better determine the possible clinical role of sortilin in the treatment of tumors with constitutively activated EGFR, this study investigated whether sortilin could also act on the nuclear EGFR signaling network.

We recently observed that EGFR–sortilin complexes were present in the nuclei of EGF-stimulated cells concomitant with genome-wide chromatin binding, with these complexes binding to transcription regulatory elements of genes associated with relapse from TKI treatment and progressive disease^12–14^. Interestingly, sortilin was found to preferentially bind to the transcription-starting site (TSS) of *cMYC*, reducing the activity of this gene. The TKI osimertinib was shown to trigger massive EGFR internalization and importation into cell nuclei of EGFR–sortilin complexes, with sortilin expression in the nuclei repressing *cMYC* expression. Because sortilin expression is significantly lower than EGFR expression in LUAD cell lines, sortilin may act as a restrictive factor, limiting EGFR transcriptional functions.

We have therefore proposed a model, in which sortilin exhibits a potential tumor-suppressor-like activity by concurrently binding to the transcription regulatory elements of EGFR-targeted genes, thereby limiting the EGFR transducing activity. The present study provides insight into the therapeutic importance of sortilin expression in LUAD, especially in EGFR-positive tumors. Sortilin may both predict the efficacy of TKIs and be a new candidate for the treatment of LUADs.

## RESULTS

### Sortilin interacts with EGFR in the nucleus

Based on findings showing that sortilin limits EGFR proliferative signaling^15,16^, we tested whether sortilin exhibits a tumor suppressor-like activity by acting on its nuclear signaling network. Although sortilin interacts physically with EGFR in A549 cells at or near the plasma membrane, as shown by red spots indicating sites of proximity ligation amplification (PLA), their interaction within the nuclei of cancer cells following EGF stimulation was not evaluated ^15,16^ (Figure 1a, insets 1-1 to 2-2). Z-stack confocal images and three-dimensional projections at 90° and 155° showed that EGFR–sortilin complexes were present in the nuclei of both EGF-stimulated and non-stimulated cells (Figure 1b, insets showing z axis #2 to #26). After incubation for 5 min, both the numbers of EGFR–sortilin clusters and their total volume in the nuclei of EGF-stimulated cells increased significantly (*p*<0.05, Figure 1c–e), suggesting that the translocation of EGFR–sortilin complexes started at early stages of EGFR endocytosis. Indeed, both immunoprecipitation (Figure 1f) and western blotting of isolated nuclei showed significant increases in EGFR–sortilin complexes (*p*<0.001, 30 min), with EGF kinetics suggesting specific sub-nuclear localizations^17^ (Figure 1g–h). Because EGFR silencing significantly reduced (*p*<0.05) the amount of sortilin in nuclear extracts despite EGF stimulation, sortilin translocation was likely not mediated by another member of the EGF family (Figure 1h and 1j).

**Figure 1:**
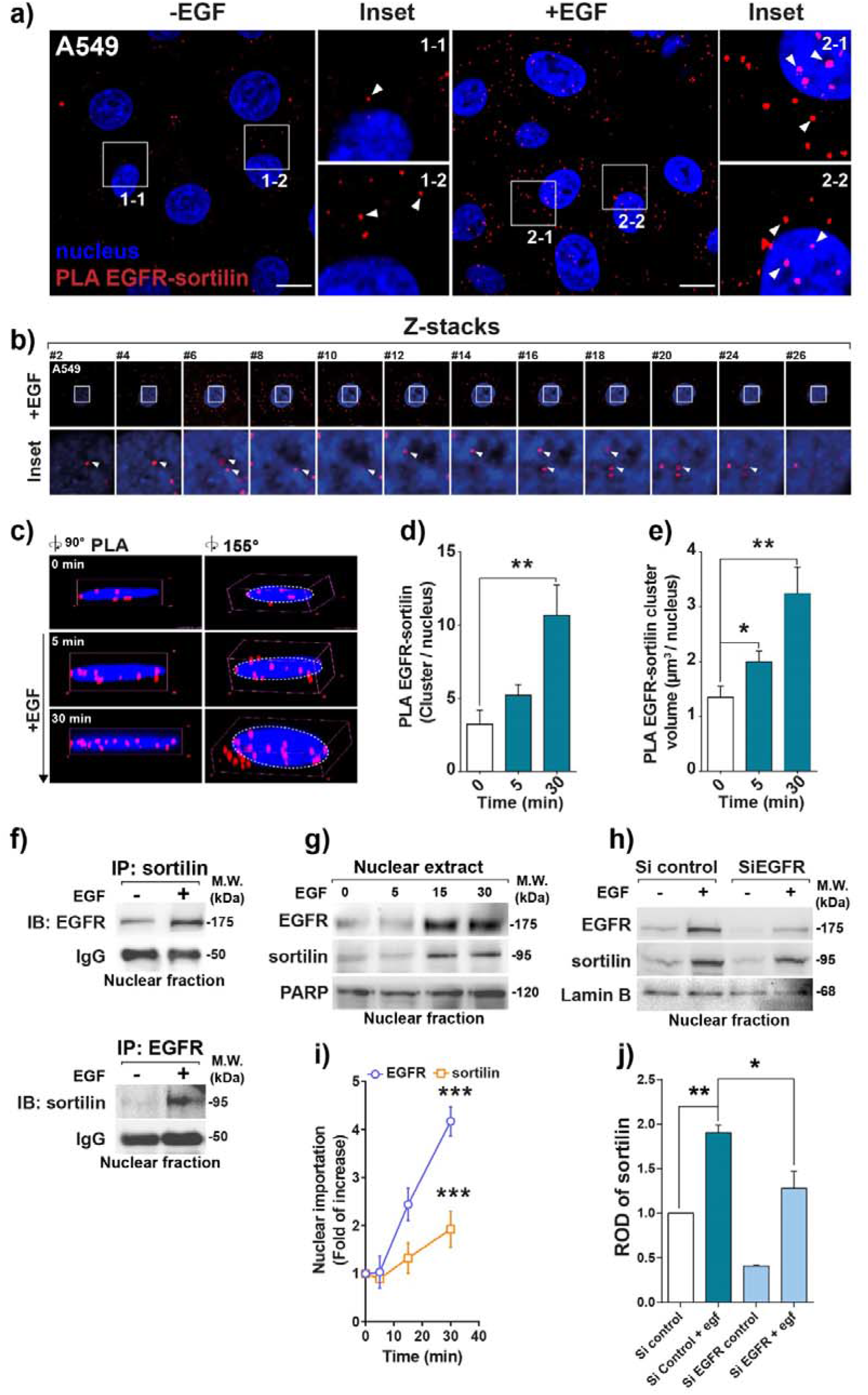
Sortilin and EGFR interact together in the nuclei of cancer cells. **(a)** Proximity ligation assay (PLA) showing the interaction between sortilin and EGFR in the lung adenocarcinoma cell line A549 in the absence or presence of EGF (50 ng/mL) for 30 min. Red spots indicate sites of PLA amplification, reflecting interactions between sortilin and EGFR. Scale bar, 10 μm; white arrows show EGFR–sortilin clusters. (**b)** Z-stack sections of confocal microscopy images showing sortilin and EGFR interactions in z axis (insets #2–26). White arrows show EGFR–sortilin clusters. **(c)** 3D confocal microscopy images showing EGFR–sortilin interactions at angles of 90° and 155°. **(d)** Quantification of EGFR–sortilin spots per nucleus, in the absence or presence of EGF for 5 or 30 min. **(e)** Estimated volumes of EGFR–sortilin clusters per nucleus (μm^3^/nucleus) in the absence or presence of EGF for 5 or 30 min. **(f)** Confirmation of EGFR–sortilin interactions by nuclear co-immunoprecipitation of A549 cell lysates in the absence or presence of EGF (50 ng/mL) for 30 min and immunoblotted (IB) with anti-EGFR antibodies. **(g)** Immunoblots showing kinetics of EGFR and sortilin nuclear importation following EGF stimulation of A549 cells. Nuclear fractions were obtained 0, 5, 15, and 30 min after stimulation with 50 ng/mL EGF. **(h)** EGFR silencing by specific siRNA transfection for 72 h before assessment of sortilin importation into the nucleus by western blotting. **(i)** Quantification of nuclear importation of EGFR and sortilin following EGF stimulation. Molecular weights (MW) are shown in kilo Daltons (kDa). **(j)** Relative optical density (ROD) of sortilin expression in isolated nuclei following EGFR depletion by siRNA. All values represent means ± SD. \**p*<0.05, \*\**p*<0.01, and \*\*\**p*<0.001 by Student’s t-test. Each experiment was repeated at least three times.

These results suggest that sortilin is imported into the nuclei of cancer cells only in the presence of EGFR, and that nuclear EGFR importation requires EGFR endocytosis. Likewise, agglomeration of EGFR–sortilin complexes in the nuclei of EGF-stimulated cells suggests a specific sub-nuclear localization that might address transcriptional functions.

### EGFR–sortilin complexes co-immunoprecipitate with chromatin

To gain insight into the role of EGFR–sortilin complexes in the nuclei of EGF-stimulated cells, we investigated whether these complexes exhibited chromatin binding properties. Chromatin immunoprecipitation assays (ChIP) were performed using micrococcal nucleases (Mnase), with the quality of enzymatic digestion validated by assessing the ability to release mono-nucleosomes (Supplementary Materials 1a-d). Because sortilin was never shown to act as a transcriptional co-factor with genomic binding sequences, and because its nuclear importation would depend on EGFR (Figure 1h and 1j), we analyzed specific DNA sequences located within the promoter regions of genes belonging to the EGF transcriptional response pathway^18^. Thus, we selected the epigenetic reprogramming gene *cMYC*^11^ and the cell cycling gene Cyclin D1 (*CCDN1*)^19^. ChIP with anti-EGFR or anti-sortilin antibodies showed that EGF stimulation resulted in the amplification of *cMYC* and *CCND1* chromatin sequences (Figure 2a). Because amplification was not observed following immunoprecipitation with their respective isotype controls (IgG1 ChIP EGFR and IgG ChIP sortilin), these results suggest that both EGFR and sortilin interact specifically with chromatin and could participate in the activity of EGF-regulated genes (Figure 2a, IgG).

**Figure 2:**
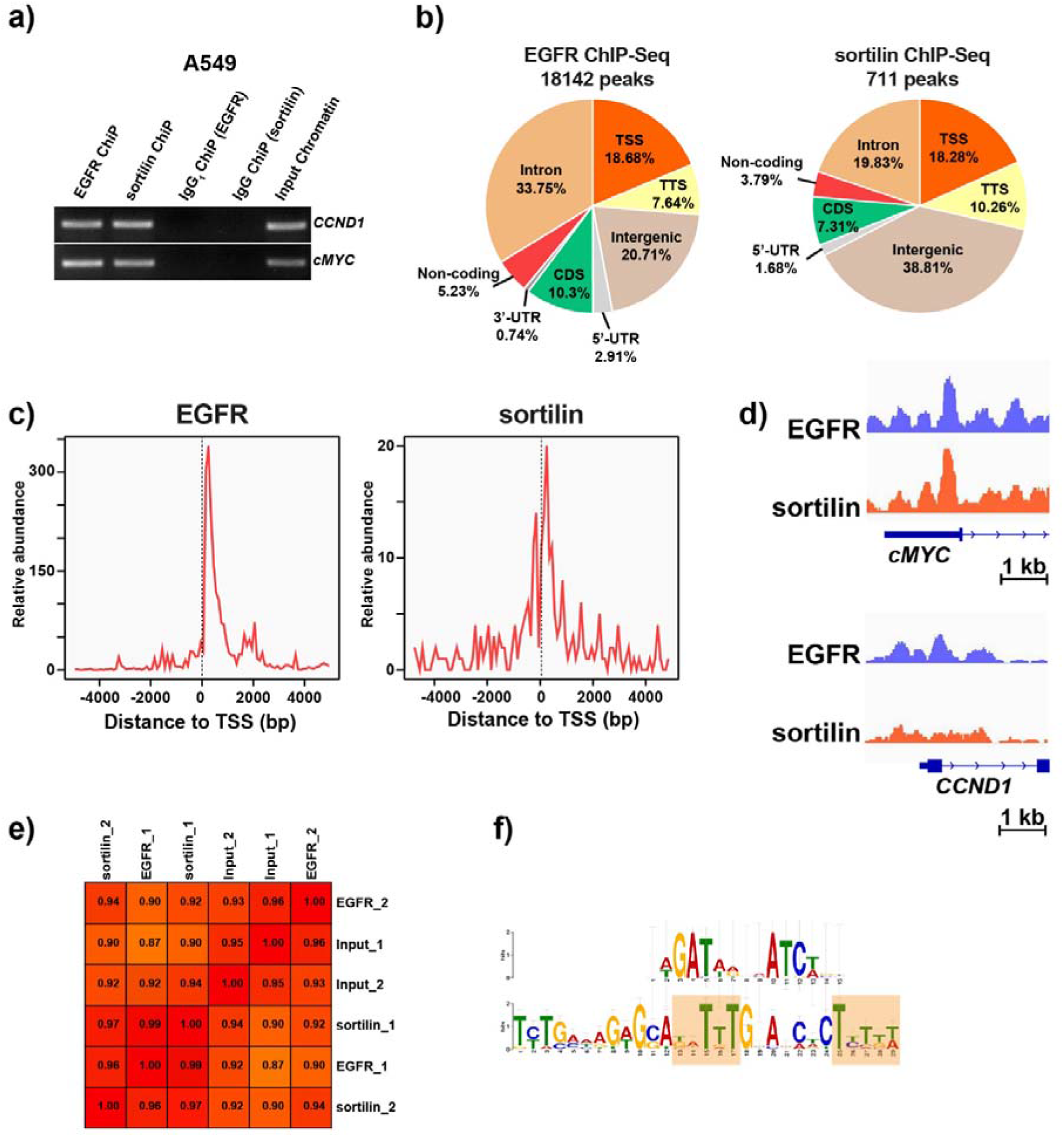
EGFR and sortilin interact with chromatin. **(a)** PCR amplification of *CCND1* and *cMYC* promoter sequences in A549 cells stimulated with EGF (50 ng/mL) for 30 min following chromatin immunoprecipitation (ChIP) with either anti-EGFR or anti-sortilin antibodies. Respective isotype IgGs, IgG1 ChIP(EGFR) and IgG ChIP (sortilin), were used as controls and compared with input samples (input chromatin) corresponding to non-ChIP DNA as internal control. **(b)** Peaks enriched for EGFR and sortilin in A549 cells stimulated with EGF (50 ng/mL) for 30 min. **(c)** Distribution of EGFR and sortilin ChIP-Seq reads near 5 kb upstream /downstream of TSS. **(d)** ChiP-Seq overview shown with the IGV genome browser representing EGFR and sortilin binding sites on *CCND1* and *cMYC* TSS. **(e)** Table showing Pearson correlation coefficients between each pair of ChIP conditions. Color intensity was representative of the magnitude of the correlation coefficient. **(f)** Common consensus sequences of EGFR and sortilin binding sites on chromatin.

To identify the DNA regions immunoprecipitated by anti-EGFR and anti-sortilin antibodies, the ChIP products were sequenced (ChIP-Seq). All libraries bound by these antibodies met all ChIP-Seq quality control criteria (Supplementary Materials 1b and c). ChIP-Seq experiments were performed on biological replicates following stimulation with 50ng/mL EGF for 30 min, with reads averaging 50 million. The percentage frequencies of peaks enriched in stimulated A549 cells were predominantly distributed within intergenic and intronic regions, as well as toward transcriptional regulating elements, including the TSS and the transcription termination site (TTS) (Figure 2b). Analysis of the segmentation of TSS sequences revealed a preferential distribution for EGFR and sortilin. ChIP-Seq peak distributions within 5 kb of TSS with aggregation plots showed that the TSS/TTS ratios for EGFR and sortilin were 2.44 and 1.78, respectively (Supplementary Figure 1). Not surprisingly, we observed an abundance of TSS peaks co-occurring with the highest expression of EGFR (Figure 2c and Supplementary Figure 1). Likewise, their overlap positions in close proximity to the TSS region suggested that EGFR–sortilin complexes affected gene activity (Supplementary Figure 1). The PLA and co-immunoprecipitation assays showing the physical interactions between EGFR and sortilin in the nuclei of A549 cells (Figure 1a–e) suggested that EGFR and sortilin have common binding sites on target loci. Significant correlations between EGFR and sortilin profiles were shown on ChiP-Seq overview using the IGV genome browser, which found EGFR and sortilin binding sites on the *CCND1* and *cMYC* TSS, as well as by Pearson’s correlation coefficients among samples (Figure 2d and 2e). Similarly, *in silico* analysis suggested that both EGFR and sortilin bound to an AT-rich minimal consensus sequence (ATRS), consisting of TNTTT or TTTNT, with N being any nucleotide (Figure 2f). These genomic sequences were previously associated with the EGFR chromatin binding site^18,19^, suggesting that EGFR–sortilin complexes potentially bind chromatin through EGFR.

Taken together with our previous results, the binding patterns of EGFR and sortilin were close to gene-proximal regulatory elements, suggesting that EGFR–sortilin complexes are involved in EGF-induced molecular processes.

### EGF stimulation enhances DNA occupancy by sortilin

To further assess whether EGF promotes EGFR and sortilin DNA binding to transcriptional regulatory elements, we designed primers corresponding to the TSS regions of genes derived from gene ontology (GO) analysis (Supplementary Table 1), followed by the use of immunoprecipitated chromatin as a qPCR template. Each immunoprecipitation met ChIP quality control (data not shown), with non-specific DNA binding ruled out by using non-relevant immunoglobulins of the same class as the respective antibodies (data not shown). A549 cells were depleted of *EGFR* and *SORT1* mRNAs ^15,16^ using specific shRNAs and incubated with antibodies to specifically immunoprecipitate chromatin. No significant differences were observed between A549 cells transfected with empty vector (pLKO cells) and wild-type A549 cells (data not shown). EGF stimulation triggered significant (*p*<0.001) chromatin binding of EGFR onto the TSS regions derived from *CCND1, cMYC*, and several genes selected by GO analysis (Figure 3a and Supplementary Figure 2a). Although EGF stimulation significantly enhanced (*p*<0.001 and *p*<0.001) sortilin binding to the TSS of selected genes (Figure 3b and Supplementary Figure 2b), amplification of the *cMYC* TSS was of especial interest. Indeed, EGF stimulation triggered a significant (*p*<0.001) reduction of sortilin chromatin binding to *cMYC* transcriptional regulatory elements when compared with control cells (Figure 3b). Because chromatin was not amplified in these mRNA-depleted cell lines (shRNA, *p*<0.0001) (Figure 3a and 3b), the differences between EGFR and sortilin binding profiles for *cMYC* TSS in basal condition may have specifically involved in *cMYC* gene activity. Likewise, because EGFR expression remains higher than that of sortilin, aggregation of free uncomplexed EGFR to sortilin could unbalance sortilin action toward gene activity. Indeed, EGFR depletion significantly (*p*<0.001) reduced the expression of *cMYC* mRNA but had no effect on *CCND1* mRNA expression (Figure 3c). By contrast, the levels of *CCND1* and *cMYC* mRNAs were significantly higher (*p*<0.001) in A549 *SORT1* mRNA-depleted than in control A549 cells (Figure 3d).

**Figure 3:**
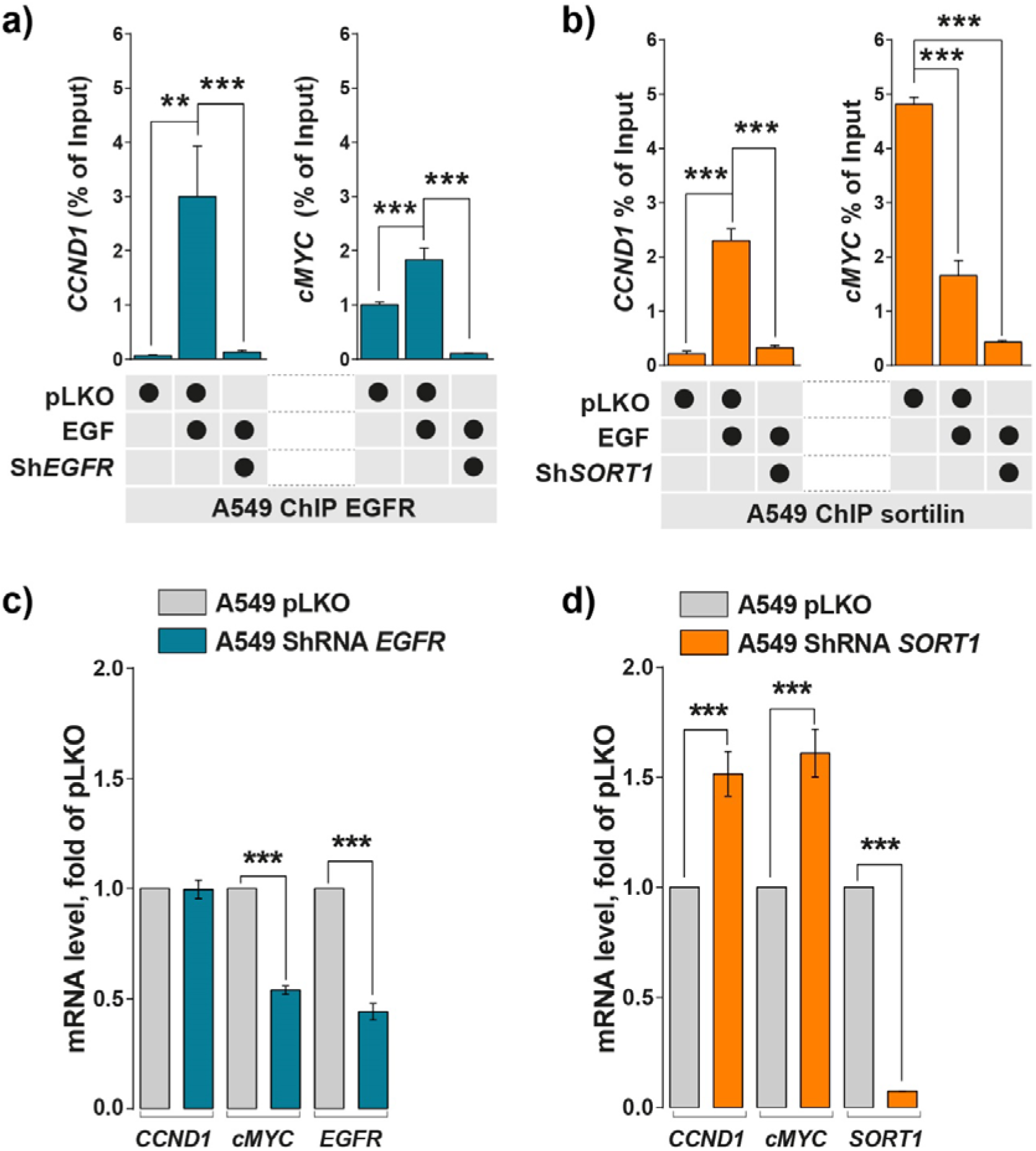
EGF stimulation increases EGFR and sortilin binding to chromatin. **(a-b)** Quantitative PCR (qPCR) of chromatin immunoprecipitated (ChIP) by anti-EGFR (blue bars) and anti-sortilin (orange bars) antibodies in control cells (pLKO) and cells depleted of *EGFR* or *SORT1* mRNA by incubation with shRNAs, and incubated in the absence or presence of EGF (50 ng/mL) for 30 min. Histograms represented the percentages of input following normalization. *CCND1* and *cMYC* promoters were amplified by qPCR. **(c-d)** RT-qPCR measurements of *CCND1* and *cMYC* mRNAs in A549 cells depleted of *EGFR* or *SORT1* mRNA with shRNAs and in control cells (pLKO). All values represent means ± SD, \*\*\**p*<0.001 by Student’s t-test. Each experiment was repeated at least three times.

Taken together, these results suggest that sortilin impairs expression of EGF response genes and could compete with EGFR at *cMYC* regulatory elements, thus limiting the expression of EGFR oncogenic co-drivers.

### Sortilin overexpression limits polymerase II recruitment to TSS

Because EGFR protein levels are decreased in H1975 cells overexpressing *SORT1* (*OE-SORT1*)^15^, we performed ChIP experiments using these cells and H1975 cells or transfected with empty vector (EV). As expected, EGF stimulation of control (EV) cells triggered significant (*p*<0.001) chromatin binding by both EGFR and endogenous sortilin (Figure 4a and Supplementary Figure 3a). By contrast, because sortilin overexpression reduced EGFR stability, EGFR chromatin binding decreased significantly despite EGF stimulation (Figure 4a and Supplementary Figure 3b). Under such experimental conditions, sortilin binding to chromatin was reduced when compared with control cells, whereas sortilin binding to the *cMYC* TSS was not altered by EGF stimulation (Figure 4b). Thus, sortilin continued to occupy the *cMYC* TSS when compared with non-stimulated *OE-SORT1* cells (Figure 4b and Supplementary Figure 3b). Using this model, we assessed the recruitment of polymerase II (Pol II), belonging to the initiating transcription complex, toward the TSS surface occupied by EGFR and sortilin (Figure 4c). Interestingly, the chromatin binding of Pol II to *cMYC* and *CCND1* TSS was significantly lower in cells overexpressing sortilin than in control cells, as was the binding of Pol II to selected genes from GO analysis (Figure 4c and Supplementary Figure 3c). Sortilin binding was higher in cells overexpressing *SORT1* than in EV cells, suggesting that sortilin impairs recruitment of Pol II and the gene activity of *CCND1* and *cMYC*. To further evaluate the consequences of increased sortilin chromatin binding, we assessed the levels in these cells of *CCND1* and *cMYC* mRNAs. Surprisingly, sortilin overexpression significantly reduced (*p*<0.001) the mRNA levels of the EGFR co-drivers *CCND1* and *cMYC* (Figure 4d).

**Figure 4:**
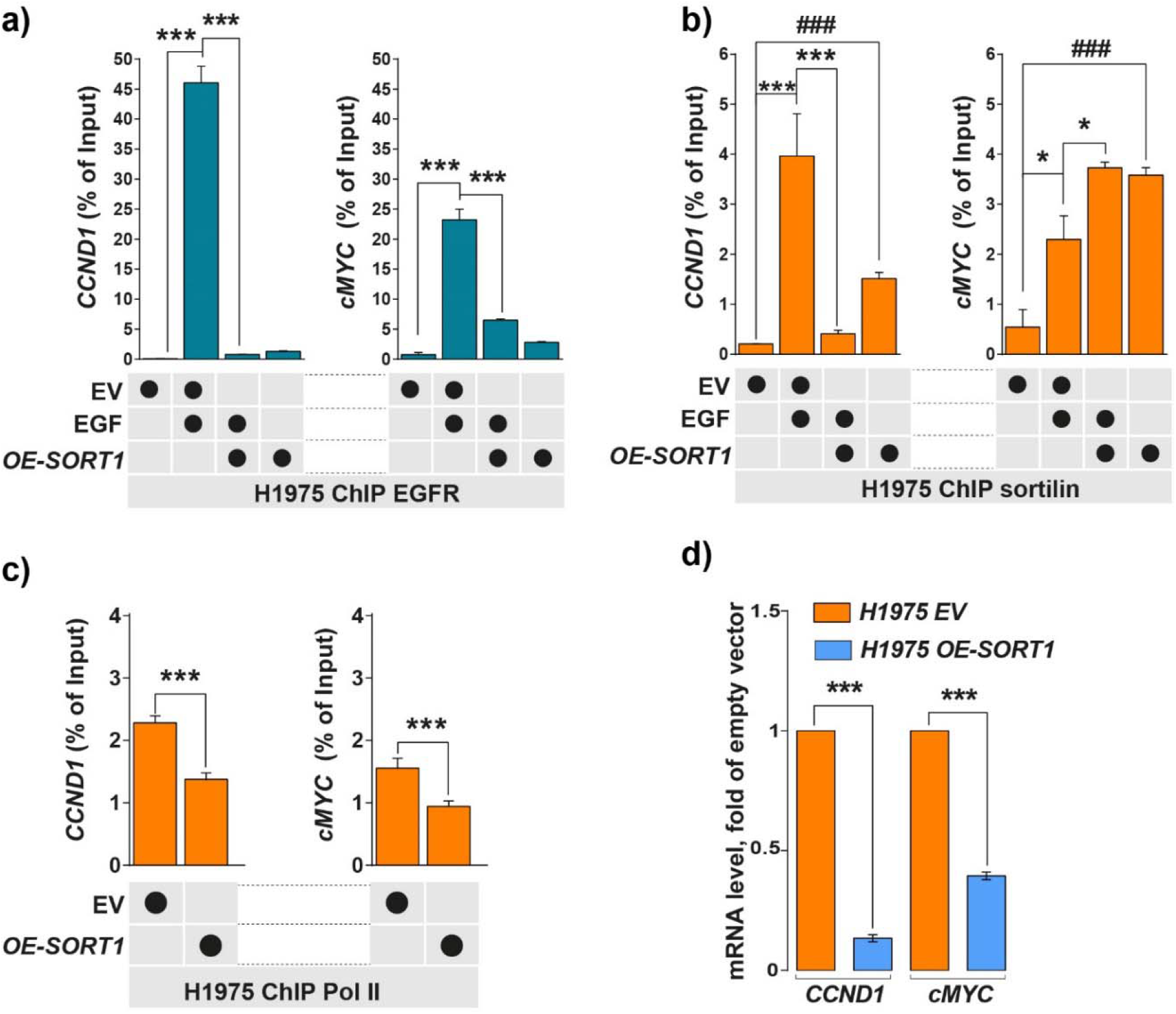
Sortilin overexpression increases sortilin chromatin binding on *cMYC* and limits polymerase II recruitment. **(a-b)** EGFR and sortilin ChIP-qPCR were performed on H1975 control cells transfected with empty vector (EV) or on H1975 sortilin overexpressing cells (*OE-SORT1*) in the absence or presence of EGF (50 ng/mL) for 30 min. *CCND1* and *cMYC* promoters were amplified by qPCR. **(c)** Pol II ChIP-qPCR performed on EV and *OE-SORT1* cells. **(d)** Levels of *CCND1* and *cMYC* mRNAs in control (EV) and *OE-SORT1* cells by qPCR. All values represent means ± SD, \**p*<0.05, \*\**p*<0.01, \*\*\**p*<0.001, and ^###^*p*<0.001 by Student’s t-tests. Each experiment was repeated at least three times.

Taken together, these results suggest that the amount of sortilin would represent a limiting factor to impair EGFR binding and Pol II recruitment at the TSS of EGF response genes. Moreover, in the presence of TKIs, the inhibition of EGFR kinase activity may result in an imbalance in sortilin chromatin binding.

### Osimertinib triggers nuclear importation of EGFR

The spatiotemporal distribution of EGFR remains critical in the treatment of patients with lung cancer, with patients relapsing due to sustained proliferative signaling in the endosome platform or enhanced nuclear importation^20^. The subcellular distribution of EGFR, however, is dependent on EGFR mutational status ^21^. For example, EGFR with a T790M mutation in H1975 cells is constitutively active, being internalized ^15,21^, whereas wild-type EGFR in A549 cells remains at the plasma membrane in the absence of ligand stimulation^15,21^. Because EGFR-targeted agents have been found to trigger EGFR endocytosis^9,22^, we investigated EGFR distribution following TKI exposure; whether inhibition of its kinase activity by osimertinib, a TKI designed to inhibit the activity of EGFR containing the T790M mutation^23^, impairs EGFR chromatin binding; and whether competition with sortilin for chromatin binding would limit the activity of this TKI. Strikingly, we found that treatment of A549 cells with 1 μM osimertinib for 24 h triggered massive EGFR endocytosis, similar to that observed by ligand stimulation with 50 ng/mL for 30 min (Figure 5a, insets 1-1 to 6-2). Cell fractionation and isolation of nuclei resulted in massive importation of EGFR in the nuclei of both cell lines, irrespective of its initial subcellular distribution (Figure 5b). Treatment with osimertinib did not inhibit EGFR importation into the nucleus, although it reduced EGFR phosphorylation. Similar to EGF stimulation (Figure 1a–f), treatment with osimertinib also resulted in the nuclear importation of sortilin, suggesting that irrespective of stimuli, sortilin could be co-imported with EGFR in a manner independent of the phosphorylation status of the latter.

**Figure 5:**
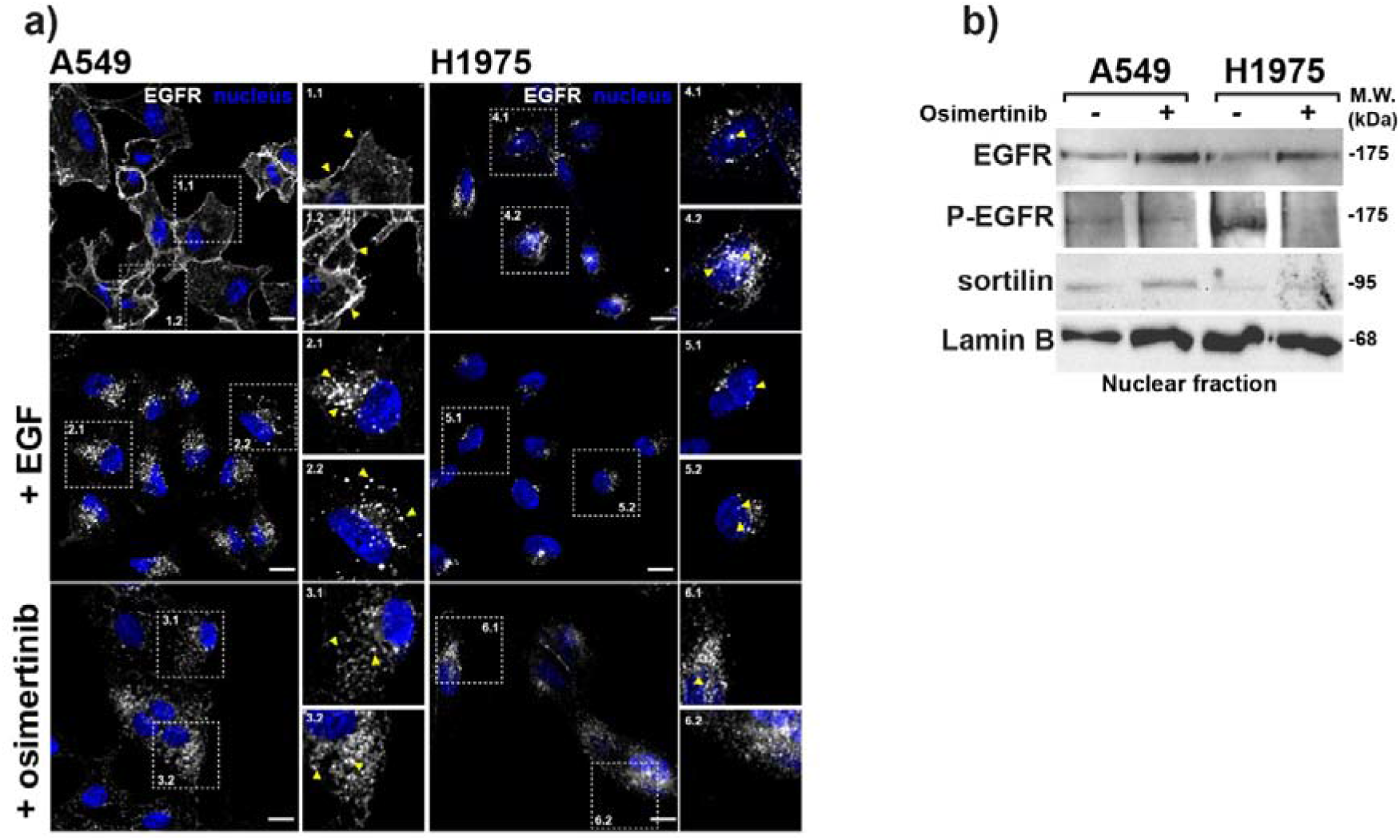
Osimertinib enhances nuclear importation of EGFR. **(a)** EGFR localization was analyzed by confocal microscopy in A549 and H1975 cells, in the absence or presence of EGF stimulation (50 ng/mL for 30 min) or osimertinib treatment (1 μM for 24 h). Scale bar, 10 μm, yellow arrows show EGFR location. **(b)** Western blotting showing that treatment of A549 and H1975 cells with osimertinib (1 μM for 24 h) controlled EGFR and sortilin importation into isolated cell nuclei following cell fractionation. Molecular weight (MW) in kilo Daltons (kDa).

### *Osimertinib increases sortilin chromatin binding to* cMYC *TSS*

We subsequently assessed whether EGFR and sortilin binding to chromatin increases following the enrichment of these proteins in the nuclear compartment. Osimertinib treatment of A549 cells carrying wild-type EGFR significantly increased EGFR binding to chromatin for each selected TSS sequence (Figure 6a and Supplementary Figure 4a). Similarly, osimertinib significantly increased sortilin binding to all chromatin sequences, except for *cMYC* TSS, where its binding remained unchanged when compared with control of A549 cells (Figure 6b and Supplementary Figure 4b). Strikingly, only *cMYC* TSS binding was significantly increased following osimertinib treatment of the EGFR-mutated H1975 cell line, whereas sortilin binding to both *CCND1* and *cMYC*, as well as to selected genes from GO analysis, increased significantly (Figure 6c and 6d, and Supplementary Figure 5a and 5b). To further analyze gene activity following chromatin binding by EGFR and sortilin, we assessed the levels of *CCND1* and *cMYC* mRNAs in these cells (Figure 6e–g). Osimertinib treatment of H1975 cells did not significantly reduce the level of *cMYC* mRNA relative to that of *CCND1* mRNA and to the level of *cMYC* mRNA in A549 cells, suggesting that EGFR and sortilin compete in binding to the *cMYC* TSS (Figure 6f). Indeed, unbalancing the proportion of sortilin in the *SORT-OE* model significantly reduced the level of *cMYC* mRNA relative to that in EV cells (Figure 6g).

**Figure 6:**
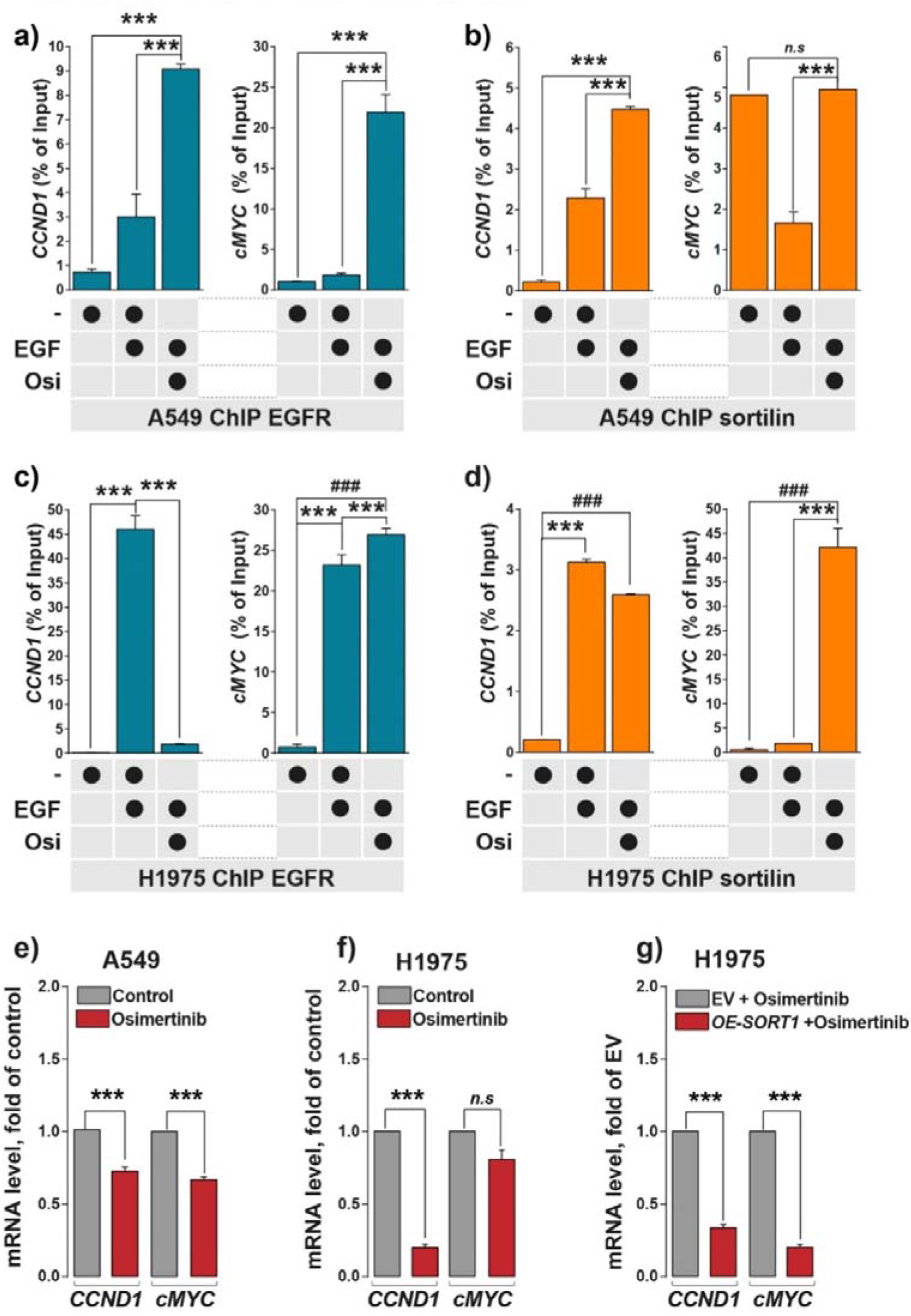
Osimertinib increases EGFR and sortilin binding to chromatin. **(a-d)** Results of EGFR and sortilin ChIP-qPCR of A549 and H1975 cells incubated in the absence or presence of EGF (50 ng/mL for 30 min) or osimertinib (1 μM for 24 h). *CCND1* and *cMYC* promoter sequences were amplified by qPCR. **(e-f)** Levels of *CCND1* and *cMYC* mRNAs determined by RT-qPCR in A549 and H1975 cells in the absence or presence of osimertinib. **(g)***CCND1* and *cMYC* mRNAs were quantified by RT-qPCR in control H1975 cells carrying empty vector (EV) and H1975 cells overexpressing (OE-SORT1) in the presence of osimertinib. All values represent means ± SD, \*\*\**p*<0.001 and ^###^*p*<0.001 by Student’s t-tests, n.s.: not significant. Each experiment was repeated at least three times.

Taken together, these results suggest that sortilin competes for binding to the regulatory elements of the *cMYC* gene, and that its expression would remain a limiting factor in the EGFR transcriptional program, irrespective of stimuli triggering its nuclear importation.

### Inverse correlation between cMYC and sortilin expression

Because uncontrolled EGFR proliferative signaling leads to cell transformation^3,24^ and ADC initiation^25–27^, and because malignant behavior is enhanced by mutation of the EGFR TK domain, we generated an inducible model (Tet-ON), in which sortilin expression was triggered in H1975 cells. Using this model, we found that treatment with doxycycline triggered sortilin expression, thereby unbalancing EGFR stability (Figure 7a). Although EGFR–sortilin complexes were increased in the nuclei of H1975^Tet-ON-*SORT1*^-induced cells, as evidenced by immunoprecipitation in the presence or absence of EGF stimulation (Figure 7b), levels of *CCND1* and *cMYC* mRNAs decreased significantly (*p*<0.001, orange histograms, Figure 7c). *In vivo*, sortilin expression triggered a significant global slowdown of tumor progression (*p*<0.001, orange curve, Figure 7d) when compared with non-induced cells (blue curve, Figure 7d). Strikingly, we also observed significant reductions in the levels of *CCND1* (*p*<0.01) and *cMYC* (*p*<0.05) mRNAs (orange histograms, Figure 7e), further suggesting that sortilin has a tumor suppressor-like activity on the expression of EGFR co-drivers associated with EGF transcriptional responses. Because these results suggested that sortilin expression would unbalance EGF transcriptional response, we assessed their clinical relevance by analyzing *SORT1* mRNA expression in 54 patients with LUAD. We found that *SORT1* mRNA levels were significantly lower (*p*<0.001) in tumor than in adjacent normal tissue samples (blue boxes, Figure 7f), findings confirmed in data sets from two other studies^28 29^ (blue boxes, Figure 7g and 7h), irrespective of disease stages (*p*<0.001) (Figure 7i).

**Figure 7:**
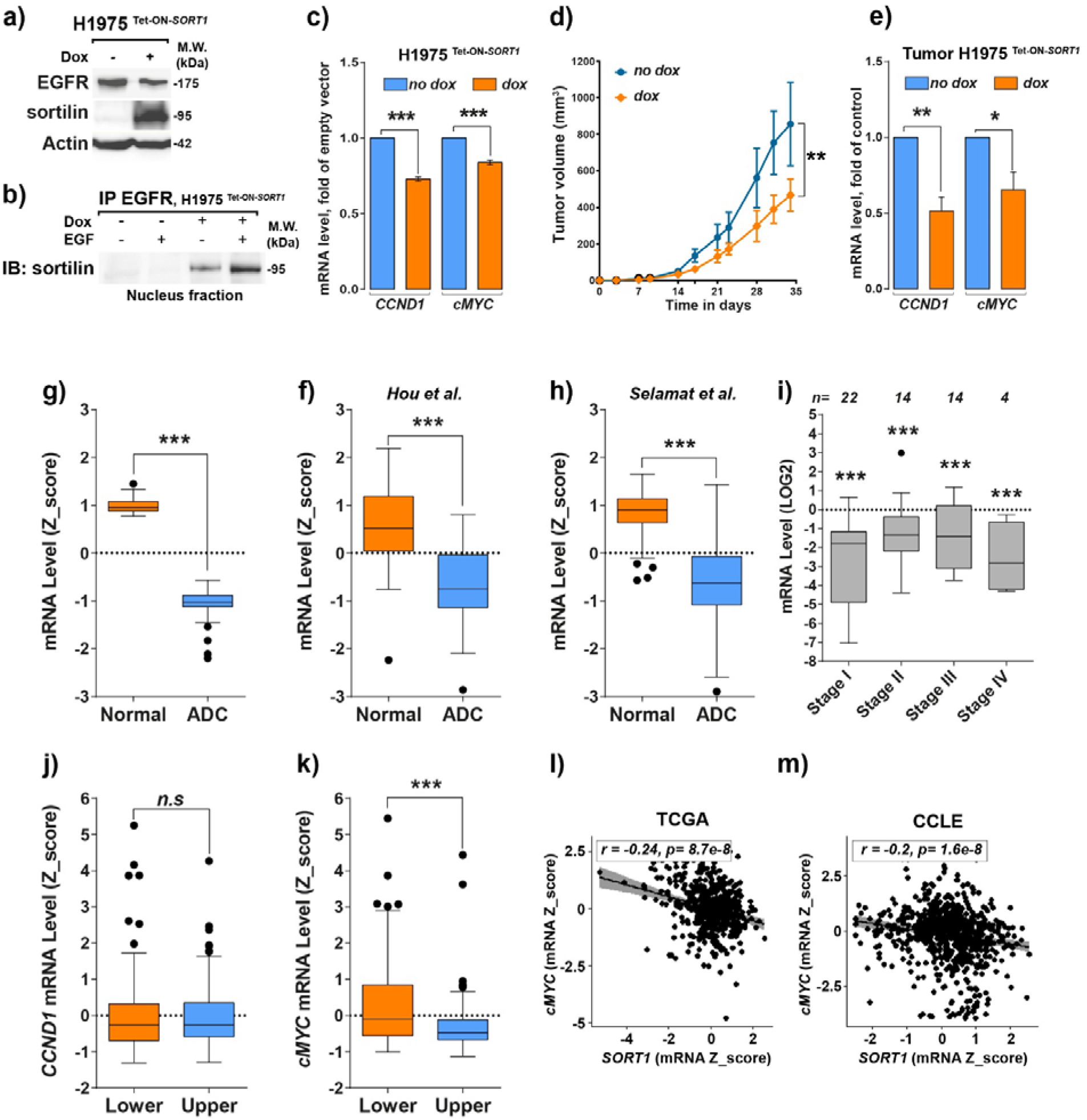
*cMYC* expression correlates inversely with *SORT1* expression *in vitro* and in tumor samples. **(a)** Western blotting showing EGFR and sortilin expression in lysates of H1975^Tet-ON-*SORT1*^ cells following incubation in the absence or presence of 100 nM doxycyclin (dox) for 24 h. **(b)** Anti-EGFR immunoprecipitation (IP) of isolated nuclei from H1975^Tet-ON-*SORT1*^ cells following incubation in the absence or presence of 100 nM doxycyclin for 24 h and stimulation with 50 ng/mL EGF for 30 min and immunoblotting (IB) with anti-sortilin. **(c)** Comparison of *CCND1* and *cMYC* mRNA levels in H1975^Tet-ON-*SORT1*^ cells following incubation in the absence or presence of 100 nM doxycycline for 24 h. **(d)** Effects of doxycyclin on tumor induction by H1975^Tet-ON-*SORT1*^ cells in NOD-SCID mice. H1975^Tet-ON-*SORT1*^ cells were subcutaneously engrafted (3×10^6^ cells/mouse) onto NOD-SCID mice. Fifteen days later, corresponding to the beginning of tumor development, mice were treated with 2 mg/mL doxycyclin in drinking water or drinking water alone, and tumor volumes were measured. Tumor growth curves are shown for mice treated with dox (orange curve) and for control mice (blue curve). **(e)** qPCR measurements of expression of *CCND1* and *cMYC* mRNAs in tumors of mice treated with (blue bar) and without (orange bar) dox. **(f-h)** Measurements of *SORT1* mRNA levels (Z-score) in normal and lung adenocarcinoma (ADC) tissue samples obtained from the **(f)** Limoges University Hospital cohort and data sets from references **(g)**28 and **(h)**29. **(i)** qPCR measurements of *SORT1* mRNA levels in tumor samples from the Limoges University Hospital cohort at different stages. **(j, k)** Quantification of **(j)***CCND1* and **(k)***cMYC* mRNA levels in tumor samples from the Limoges University Hospital cohort expressing the lowest and highest quartiles of sortilin expression. **(l)** Correlation between levels of *cMYC* and *SORT1* mRNA levels in NSCLC patients in the TCGA database (*r*=-0.24; *p*=8.7.10^-8^) and **(m)** in solid cancer cell lines from the Cancer Cell Line Encyclopedia (CCLE) database (*r*=-0.2; *p*=1.6.10^-8^). Diagrams represent the correlation between *SORT1* expression and *cMYC* expression. All values are expressed as means ± SD, \*\**p*<0.01 and \*\*\**p*<0.001 by Student’s t-test, n.s.: not significant. Each experiment was repeated at least three times.

We therefore categorized these patients by quartiles of *SORT1* mRNA expression and compared their levels of expression of other mRNAs. Interestingly, we found that only *cMYC* mRNA expression was affected by the level of *SORT1* mRNA expression, with *cMYC* mRNA expression being significantly lower (*p*<0.001, blue box, Figure 7k) in patients with high sortilin expression (Upper). We also evaluated the effects of sortilin expression on *cMYC* expression in several publicly available data sets from the MSKCC cBioPortal^30,31^, including 240 patients in The Cancer Genome Atlas (TCGA)^32^ and 665 solid cancer cell lines in the Cancer Cell Line Encyclopedia (CCLE)^33^. Strikingly, c*MYC* expression was inversely correlated with *SORT1* expression in both patient tissue samples (*r*=-0.24, *p*=8.7.10^-8^) and cancer cell lines (*r*=-0.2, *p*=1.6.10^-8^).

Taken together, these findings suggest that sortilin alters the activity of the epigenetic reprogramming gene, *cMYC*. Because sortilin remains dysregulated in malignant tissues, enabling an imbalance in the EGF transcriptional response, the malignant behavior of tumors with mutant EGFR would be increased by the expression of co-oncogenic drivers despite the presence of a TKI.

## DISCUSSION

The present study showed that sortilin is a key regulator of nuclear EGFR and that it limits EGFR transducing activity. These findings suggest a mechanism for the biological activity of sortilin. In this model, sortilin interacts with EGFR at the chromatin regulatory elements of EGF response genes, such as those involved in cell reprogramming (*cMYC*) and proliferation (*CCND1*), both of which are hallmarks of cancer, with sortilin limiting their expression ^34^. Figure 8 summarizes the role of sortilin in nuclear EGFR networking and its putative underlying mechanism. We had previously shown that sortilin plays an important striking role in directing EGFR toward rapid internalization and degradation following EGF stimulation^15^. PLA and nuclear IP immunoprecipitation experiments in the present study showed the spatiotemporal distribution of EGFR–sortilin complexes in the nuclei of EGF-stimulated cells. Chromatin immunoprecipitation and genome-wide analysis revealed that EGFR and sortilin were coordinately organized in complexes directed toward the regulatory elements of EGF response genes^35,36^. Indeed, the loci co-occupied by EGFR–sortilin were similar to those revealed by transcriptomic gene expression and genome-wide analysis^35,36^. The preferential accumulation of these complexes at TSS containing the EGFR binding chromatin sequence ATRS^18,37^ suggest that they bind to chromatin through EGFR.

**Figure 8:**
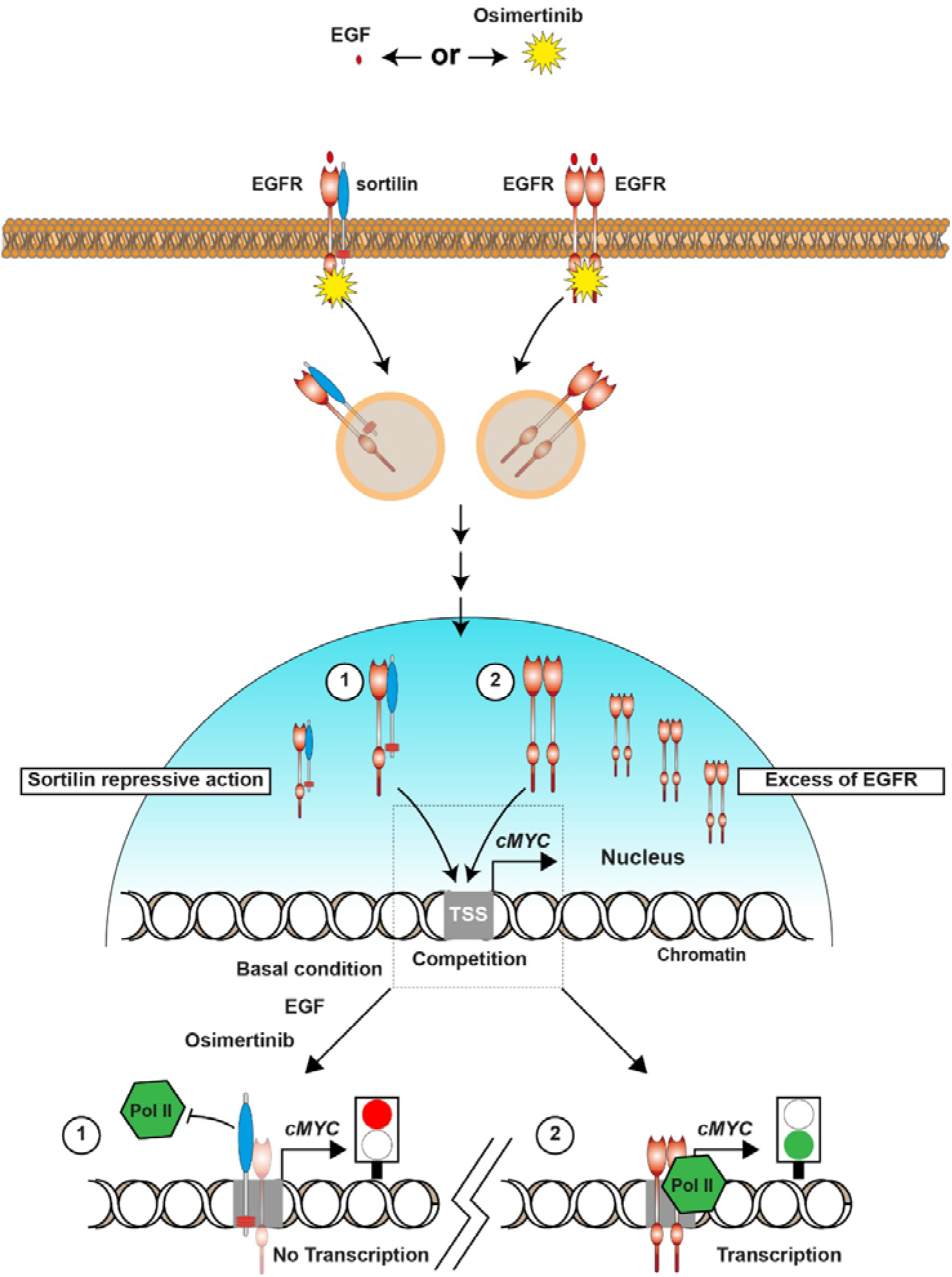
Model of sortilin regulation of transcription. Schematic diagram showing that sortilin has tumor suppressor-like activity, reducing co-oncogene transcription. EGF activates EGFR and induces its internalization as a homodimer or as a hetero dimer with sortilin. Osimertinib treatment promotes EGFR internalization and nuclear translocation. (1) Endocytosis of EGFR with sortilin can result in translocation of the complex into the nucleus, where it binds to chromatin at the TSS, thereby repressing RNA Pol II binding and cMyc co-oncogene transcription. (2) Excess EGFR homodimers imported into the nucleus bind to a specific chromatin area and trigger the recruitment of RNA Pol II, activating transcription.

Because the expression of sortilin in NSCLC cell lines is low^15^, the role of sortilin in the EGFR transcriptional program was delineated using both constitutive and inducible models of *SORT1* expression in the highly aggressive cell line H1975, which expresses EGFR carrying the mutation T790M. Although sortilin affected EGFR stability, their interaction in the nucleus increased, as did sortilin chromatin binding. In this model, both EGFR and Pol II binding to the TSS surface of *cMYC* and *CCND1* decreased significantly, as did the levels of their respective mRNAs. Likewise, *SORT1* expression *in vivo* triggered a global slowdown of tumor progression, along with significant reductions in the levels of *cMYC* and *CCND1* mRNAs. These results suggest that sortilin was able to bind TSS sequences irrespective of stimuli, and that sortilin expression remains also crucial to limit EGFR nuclear networking.

These observations raised questions concerning whether neo-endocytosed EGFR could result in imbalances in nuclear EGFR–sortilin complexes. We therefore treated cells with the TKI osimertinib, which inhibits the kinase activity of EGFR, thereby limiting its phosphorylation and endocytosis, the first step in its nuclear importation. Although EGFR phosphorylation decreased, both EGFR and sortilin were imported into cell nuclei, increasing their binding to chromatin in A549 cells bearing wild-type EGFR, and markedly increasing *cMYC* mRNA in H1975 cells. Because, *cMYC* expression in osimertinib-treated H1975 cells decreased significantly only when sortilin was overexpressed, the amount of sortilin may be insufficient to alleviate the EGFR transcriptional program, particularly regarding *cMYC* gene activity. Sortilin expression was found to decrease with the pathologic grade of tumors^15^, consistent with findings in this study showing that sortilin is downregulated in most malignant tissues. An assay of tissue samples from 54 patients with LUAD showed that only *cMYC* mRNA level was significantly decreased in malignant tissues with high levels of *SORT1* mRNA. A similar inverse correlation in *cMYC* and *SORT1* expression was observed in tumor tissues and solid cancer cell lines in publicly available datasets. Taken together, these results provide new insights into the tumor suppressor-like activity of sortilin, showing that it alters *cMYC* gene activity. Interestingly, *cMYC* belongs to the panel of genes co-occurring with the EGFR T790M mutation^13^. Because *cMYC* expression reprograms cells, resulting in the formation and maintenance of tumor-initiating cells endowed with metastatic capacities^11^, these cells become resistant to both anti-EGFR therapy^12^ and radiotherapy^38^.

In summary, our findings provide insight into the role of sortilin in LUAD. Sortilin binds to the chromatin elements of EGF response genes, thereby repressing *cMYC* transcription. This potential mechanism of regulation suggests that sortilin expression may be predictive of tumor responses to anti-EGFR treatment and patient outcomes.

## MATERIALS & METHODS

### Chromatin immunoprecipitation (ChIP) assay

Chromatin immunoprecipitation assays were performed using SimpleChIP® Enzymatic Chromatin IP Kits (Magnetic Beads) (#9003, Cell Signaling, Ozyme, France). Briefly, about 2.10^7^ cells were crosslinked with 1% formaldehyde for 10 min at room temperature. The formaldehyde reaction was quenched by adding glycine solution (#7005, Cell Signaling), followed by incubation for 5 min at room temperature. Crosslinked cells were harvested by centrifugation at 2000 x g for 5 min, washed twice with 20 mL ice-cold phosphate buffered saline (PBS, Gibco, France), and again centrifuged. Each cell pellet was resuspended in 4 mL of 1X Nuclei isolation buffer A (#7006, Cell Signaling) containing 1 M dithiothreitol (DTT) (#7016, Cell Signaling) and protease inhibitor cocktail (PIC) (#7012, Cell Signaling), followed by incubation for 10 min on ice and centrifugation at 2000 x g for 6 min at 4°C. Each pellet was resuspended in 4 mL of 1X Nuclei isolation buffer B (#7007, Cell Signaling) supplemented with 1 M DTT, centrifuged at 2000 x g for 5 min at 4°C, resuspended in 400 μL buffer B containing 2 μL Micrococcal Nuclease (#10011, Cell Signaling), and incubated for 20 min at 37°C with frequent mixing. DNA digestion was stopped by adding 0,5 M EDTA (#7011, Cell Signaling) and incubating on ice for 2 min. Nuclei were harvested by centrifugation at 16,000 x g for 1 min at 4°C, resuspended in 1X ChIP buffer (#7008, Cell Signaling) containing PIC, and lysed by sonification, and the lysates were centrifuged at 9 400 x g for 10 min at 4°C. Following purification from the supernatant, the sizes and concentrations of DNA fragments were evaluated by 2% agarose electrophoresis and NanoDrop™ quantification (NanoDrop™ ND2000C, Thermo Scientific™, France). Immunoprecipitation assays were performed by mixing 50 μg DNA, 500 μL of 1X ChIP buffer with PIC, and 2 μg antibody to EGFR H11 (anti-EGFR H11, #MA5-13070, ThermoFisher Scientific™, France), sortilin (#ANT-009, Alomone, Israël), normal Rabbit IgG (#2729, Cell Signaling), or mouse (G3A1) mAb IgG1 isotype control (#5415S, Cell Signaling). The mixtures were incubated overnight at 4°C with rotation, and 30 μL ChIP-Grade Protein G Magnetic Beads (#9006, Cell Signaling) were added, followed by incubation for 3 h at 4°C with rotation. The beads were washed three times with low salt wash buffer (1X ChIP buffer) and once with high salt buffer (1X ChIP buffer; 1M NaCl). DNA and proteins were eluted from beads by adding 150 μL of 1X elution buffer (#7009, Cell Signaling) and heating at 65°C for 30 min. Supernatants were harvested and digested by adding 2 μL proteinase K (#10012, Cell Signaling) and incubating overnight at 65°C. DNA was purified by loading onto Purification Columns (#10010, Cell Signaling) and eluting in 40 μL DNA elution buffer (#10009, Cell Signaling). ChIP assays were performed by qPCR using the fold enrichment method, which was based on differences in DNA quantity between specific antibody conditions and isotypic conditions of immunoprecipitation.

### Subcellular fractionation

Nuclear and cytoplasmic fractions were extracted from cells using NE-PER™ Nuclear and Cytoplasmic Extraction Reagent kits (Thermo Scientific™). Briefly, about 1.10^6^ cells were harvested with trypsin-EDTA and centrifuged at 500 x g for 5 min. The cell pellets were washed with ice-cold PBS (Gibco) and harvested by centrifugation at 500 x g for 5 min. The cells were resuspended in Cytoplasmic Extraction Reagent I (CER I), mixed, and incubated on ice for 10 min. Cytoplasmic Extraction Reagent II (CER II) was added to the cell suspensions, which were incubated for 1 min on ice and centrifuged at 16 000 x g for 5 min at 4°C. The cytoplasmic fractions were harvested, and the pellets were washed with PBS and re-centrifuged. These nuclear pellets were resuspended in Nuclear Extraction Reagent (NER) and incubated on ice for 40 min, with mixing every 10 min. These nuclear lysates were centrifuged at 16 000 x g for 10 min at 4°C, and the nuclear fractions were harvested immediately. Subcellular fractionation was evaluated by western blotting. During these extractions, the CER I: CER II: NER volume ratios were maintained at 200: 11: 100 μL.

### Nuclear immunoprecipitation

Following the extraction of nuclear fractions, nuclear immunoprecipitations were performed using NE-PER™ Nuclear and Cytoplasmic Extraction Reagent kits (Thermo Scientific™). Briefly, nuclear extracts were diluted with radioimmunoprecipitation assay (RIPA) buffer (50 mM Tris-HCl pH 8.0, 150 mM NaCl, 1% Nonidet P-40 (NP-40), 5% sodium deoxycholate, 0.1% sodium dodecyl sulphate (SDS), 1 mM sodium orthovanadate, 1 mM NaF, 1% protease inhibitors). Antibodies for immunoprecipitation were incubated with Dynabeads™ linked to Protein G (#10003D, Invitrogen™) for 10 min at room temperature. Nuclear lysates were added, followed by incubation for 2 h at room temperature with agitation. The beads were washed three times with PBS (Gibco), and bound proteins were eluted by incubation with 2X Laemmli loading buffer (4% SDS, 10% 2-mercaptoethanol, 20% glycerol, 0.004% bromophenol blue, 0.125 M Tris-HCl) at 95°C for 10 min. SDS-PAGE and western blotting analysis were subsequently performed.

### Immunoblotting

Cells were washed with ice-cold PBS (Gibco) and lysed with cell lysis buffer (4% SDS, 10% 2-mercaptoethanol, 20% glycerol, 0.125 M Tris-HCl pH 6.8) containing 1% PIC (#7012, Cell Signaling). The cell lysates were sonicated on a Vibra-Cell Sonifier, set at 60% amplitude, three times for 5 sec each, with at least 1 min on ice between pulses. The lysates were centrifuged at 16 000 x g for 20 min at 4°C, and their protein concentrations were measured by Bradford protein assays. Aliquots containing 40 μg protein were loaded onto SDS-PAGE gels, with western blot analysis performed using specific antibodies against sortilin (#Ab16640, Abcam, France), P-EGFR (Tyr 1068, #3777, 1:1000 dilution; Cell Signaling), EGFR (#4267, 1:1000 dilution, Cell Signaling; clone H11 #MA5-13070, 1:500 dilution, Fisher Scientific, France), pERK1/2 (Thr202/Thr204, #4370, 1:1000 dilution, Cell Signaling), ERK1/2 (#9102, 1:1000 dilution, Cell Signaling), pAKT (Ser 473, #4060, 1:1000 dilution, Cell Signaling), AKT (#4691, 1:1000 dilution, Cell Signaling), lamin b1 (#HPA050524, 1:1000 dilution, Atlas Antibodies), tubulin (#sc-53646, Santa Cruz Biotechnology, Tebu, France), and actin (#A2066, 1:10000 dilution, Sigma, France), with the latter used as a loading control. The blots were subsequently incubated with horseradish peroxidase (HRP)-conjugated secondary antibodies (Dako, 1:1000 dilution, Agilent, France) and enhanced chemiluminescence substrate.

### Cell culture

The A549 and H1975 cell lines were obtained from the American Type Culture Collection (ATCC) and cultured in Dulbecco’s modified Eagle’s medium GlutaMAX (Gibco) supplemented with 10% fetal bovine serum (IDbio, France), 1% antibiotics (Gibco), and 1% non-essential amino acids (Gibco) at 37°C in a humidified atmosphere containing 5% CO_2_. Where indicated, cells were stimulated with 50 ng/mL EGF for 30 min, or treated with 1 μM of the TKI Osimertinib (AZD9291, Tagrisso, Cliniscience, France) for 24 h.

### Mice and *in vivo* tumor growth

Female NOD-SCID mice obtained from Janvier Labs (France) were housed in a control non-pathogen atmosphere. All experiments were performed in accordance with the French Veterinary Department. About 1.10^6^ H1975 cells overexpressing sortilin in the presence of doxycycline were engrafted onto the left thigh of each mouse. Tumor volume, calculated as length×width×(length+width)/2, was measured twice weekly. Following tumor development, mice were or were not administered 2 mg/mL doxycycline in drinking water. The mice were sacrificed 34 days after cell engraftment, and their tumors were collected. One part of each tumor was fixed in formaldehyde and embedded in paraffin for immunohistochemistry, whereas a second part was used to assess mRNA and protein overexpression by qPCR and western blotting, respectively.

### Immunofluorescence and confocal microscopy

Cells grown on glass coverslips were washed twice in ice-cold PBS before fixation in methanol or 4% paraformaldehyde for 10 min on ice. The cells were washed with PBS containing 1% (w/v) BSA (IDbio) and incubated for 30 min with PBS containing 3% BSA. The cells were immunolabeled at 4°C overnight with primary antibody to EGFR (Cell Signaling, Ozyme, #4267) or sortilin (Abcam, #ab16640, France), each diluted 1:100 in blocking solution. The cells were subsequently washed three times with PBS containing 1% BSA, incubated with Alexa Fluor 594-conjugated anti-rabbit IgG or Alexa Fluor 488-conjugated anti-mouse IgG antibodies (1:1000; Life Technologies, France) for 2 h at room temperature, and again washed three times with PBS containing 1% BSA. The cells were mounted using Fluoroshield mounting medium (Sigma), containing 4, 6-diamidino-2-phenylindole (DAPI) to stain the nuclei. Endocytic assays were performed using biotinylated EGF complexed to Alexa Fluor 647, according to the manufacturer’s instructions (Life Technologies, #E35351). Fluorescent images were obtained using epifluorescence microscopes (Zeiss Axiovert), equipped with a laser-scanning confocal imaging system (Zeiss LSM 510 META or LSM800). Mander’s coefficients were calculated using the Zeiss LSM 510 META or ZEN software (Zeiss) on non-saturated pictures with optical slices of 0.8 μm. At least 30 cells were acquired for each condition. Cell surface expression of EGFR and sortilin, each calculated from the difference between the whole-cell and intracellular means of fluorescence, were analyzed using ImageJ software (NIH). For PLA, the cells were fixed with 4% paraformaldehyde for 10 min, permeabilized in PBS containing 0.1% Triton X-100 (Sigma) for 30 min on ice, and washed with PBS. The cells were subsequently incubated in blocking solution (2% BSA in PBS) for 30 min at 37°C in a humidified chamber, followed by incubation with primary antibodies against EGFR (mouse monoclonal, Life Technologies) and sortilin (rabbit polyclonal, Abcam), each diluted 1:100 in blocking solution, for 30 min at 37°C. The cells were washed with buffer A from the Duolink II proximity ligation assay kit (Olink Bioscience, Sigma), followed by the addition of Duolink II PLA probe anti-mouse Minus and Duolink II PLA probe anti-rabbit Plus, and incubation for 60 min at 37°C. To link the two probes, the cells were washed in buffer A and incubated for 30 min at 37°C in Duolink II ligation buffer diluted in filtered distilled water containing ligase. Following ligation, the cells were washed in buffer A and incubated for 100 min at 37°C with the Duolink II orange amplification buffer containing polymerase. The cells were then washed three times in buffer B and mounted with in-situ mounting medium containing DAPI. Quantitative analyses of each independent sample were performed using ImageJ software (NIH, Bethesda, Maryland, USA), based on the mean fluorescence values. At least 50 cells were acquired for each condition, with the results presented as ratios relative to control cells.

### Plasmids and lentivirus-mediated RNA interference

The JetPei transfection reagent (Polyplus Transfection, Ozyme, France) was utilized for both transient and stable transfection of cells. Inducible sortilin overexpressing cell lines were generated by lentivirus-mediated RNA interference. Briefly, H1975 cells were infected twice, once with lentivirus containing DNA encoding a Tet-On system and then with lentivirus encoding sortilin overexpression. About 5×10^5^ cells were infected in complete medium containing 8 μg/mL polybrene (Sigma) and concentrated lentivirus (five lentiviral particles/cell) for 48 h, followed by selection with blasticidine (1 μg/mL, Sigma). The cells were subsequently re-infected with the second type of lentivirus before selection with puromycin (1 μg/mL, Sigma).

### Total RNA extraction and quantitative (q-)PCR analysis

Total RNA was extracted from 50 mg tissue or about 1.10^6^ cells using QIAzol Lysis Reagent (#79306, QIAGEN, France). Briefly, tissues or cells were lysed in QIAzol reagent before the addition of chloroform and centrifugation. The aqueous phase of each sample was decanted, followed by precipitation with isopropanol at −80°C for 1 h and centrifugation at 16 000 x g for 10 min at 4°C. The RNA pellets were washed with 75% ethanol, again centrifuged at 16 000 x g for 10 min at 4°C, and resuspended in water. Aliquots containing 2 μg total RNA were reverse transcribed to cDNA using Superscript III (Invitrogen), according to the manufacturer’s protocol. Each qPCR reaction contained 50 ng cDNA, TaqMan probes specific to each mRNA (Table), and Premix Ex Taq (#RR39WR, TaKaRa, France), with amplifications performed on a QuantStudio 3 real-time thermal cycler (Applied Biosystems, France). The results of RT-qPCR for each gene were normalized to those of ACTB mRNA expression in the same samples using the ΔΔCt method. ChIP-qPCR probes were designed to be complementary to the genomic DNA promoter sequence of each targeted gene and were synthesized by the custom TaqMan service from ThermoFisher Scientific.

**Table:** Probes synthesized for RT-qPCR

| Targeted gene | TaqMan™ probes references |
| --- | --- |
| ACTB | Hs01060665_g1 |
| CCND1 | Hs00765553_m1 |
| DHODH | Hs00361406_m1 |
| DUSP12 | Hs00170898_m1 |
| EGFR | Hs01076090_m1 |
| MYC | Hs00153408_m1 |
| SNAPC1 | Hs00608182_m1 |
| SORT1 | Hs00361760_m1 |
| STX6 | Hs01057343_m1 |

### Statistical analysis

Relative fluorescence intensities and the results of western blotting and ChIP experiments were compared with controls using PAST software (version 2.17). Data shown are representative of at least three independent experiments. Error bars represent the standard error of the mean. Results were analyzed for statistical significance by ANOVA, with 0.05 considered statistically significant. Correlations between levels of *cMYC* and *SORT1* mRNAs in the TCGA and CCLE databases were evaluated by linear regression analysis using R software (version 3.6.1).

## Acknowledgements

This study was generously supported by Chaire de Pneumologie Expérimentale from Association Limousine d’Aide aux Insuffisants Respiratoires-Assistance Ventilatoire à Domicile (ALAIR-AVD; Limoges, France), the Foundation of the University of Limoges, the Comité d’Orientation de la Recherche sur le Cancer en Limousin, and the Ligue Contre le Cancer. E.L. was supported by a doctoral fellowship from the Association du Développement Education Recherche-Limousin Poitou-Charentes (ADER-LPC). The authors thank all their colleagues who contributed their time and materials to this study, and thank Nicolas Vedrenne for his assistance with experiments on mice and human tissue samples. The authors are especially grateful to Alain Chaunavel from the “Centre de Ressources Biologiques Biolim,” Department of Pathology, University Hospital Limoges, and Claire Carrion from the Imaging Cytometry Platform of the University of Limoges, for technical support.

## Author contributions

L.E. and G.C. performed the experiments and analyzed the data. T.J. and C.A participated in the collection of patient samples and clinical data. G.F., J. M-O., B.F., M.B., V.F., N.T. and L.F. participated in the study design. G.F., J.M-O., F.V., N.T. and L.F. coordinated the study. All authors have read and approved the final manuscript.

## Data availability

All relevant data are available from the corresponding authors on request.

## References

1. Herbst, R. S., Heymach, J. V. & Lippman, S. M. Lung cancer. N. Engl. J. Med. 359, 1367–1380 (2008).

2. Sharma, S. V., Bell, D. W., Settleman, J. & Haber, D. A. Epidermal growth factor receptor mutations in lung cancer. Nat. Rev. Cancer 7, 169 (2007).

3. Di Fiore, P. P. et al. Overexpression of the human EGF receptor confers an EGF-dependent transformed phenotype to NIH 3T3 cells. Cell 51, 1063–1070 (1987).

4. McDermott, U. et al. Identification of genotype-correlated sensitivity to selective kinase inhibitors by using high-throughput tumor cell line profiling. Proc. Natl. Acad. Sci. U. S. A. 104, 19936–19941 (2007).

5. Rosell, R. et al. Erlotinib versus standard chemotherapy as first-line treatment for European patients with advanced EGFR mutation-positive non-small-cell lung cancer (EURTAC): a multicentre, open-label, randomised phase 3 trial. Lancet Oncol. 13, 239–246 (2012).

6. Pi, C. et al. EGFR mutations in early-stage and advanced-stage lung adenocarcinoma: Analysis based on large-scale data from China. Thorac. Cancer 9, 814–819 (2018).

7. Chong, C. R. & Jänne, P. A. The quest to overcome resistance to EGFR-targeted therapies in cancer. Nat. Med. 19, 1389–1400 (2013).

8. Liccardi, G., Hartley, J. A. & Hochhauser, D. EGFR nuclear translocation modulates DNA repair following cisplatin and ionizing radiation treatment. Cancer Res. 71, 1103–1114 (2011).

9. Li, C., Iida, M., Dunn, E. F., Ghia, A. J. & Wheeler, D. L. Nuclear EGFR contributes to acquired resistance to cetuximab. Oncogene 28, 3801–3813 (2009).

10. Hsu, S.-C. & Hung, M.-C. Characterization of a novel tripartite nuclear localization sequence in the EGFR family. J. Biol. Chem. 282, 10432–10440 (2007).

11. Poli, V. et al. MYC-driven epigenetic reprogramming favors the onset of tumorigenesis by inducing a stem cell-like state. Nat. Commun. 9, 1024 (2018).

12. Strippoli, A. et al. c-MYC Expression Is a Possible Keystone in the Colorectal Cancer Resistance to EGFR Inhibitors. Cancers 12, (2020).

13. Blakely, C. M. et al. Evolution and clinical impact of co-occurring genetic alterations in advanced-stage EGFR-mutant lung cancers. Nat. Genet. 49, 1693–1704 (2017).

14. Santoni-Rugiu, E. et al. Intrinsic resistance to EGFR-Tyrosine Kinase Inhibitors in EGFR-Mutant Non-Small Cell Lung Cancer: Differences and Similarities with Acquired Resistance. Cancers 11, (2019).

15. Al-Akhrass, H. et al. Sortilin limits EGFR signaling by promoting its internalization in lung cancer. Nat. Commun. 8, 1182 (2017).

16. Wilson, C. M. et al. Sortilin mediates the release and transfer of exosomes in concert with two tyrosine kinase receptors. J. Cell Sci. 127, 3983–3997 (2014).

17. Faraco, C. C. F. et al. Translocation of Epidermal Growth Factor (EGF) to the nucleus has distinct kinetics between adipose tissue-derived mesenchymal stem cells and a mesenchymal cancer cell lineage. J. Struct. Biol. 202, 61–69 (2018).

18. Lin, S. Y. et al. Nuclear localization of EGF receptor and its potential new role as a transcription factor. Nat. Cell Biol. 3, 802–808 (2001).

19. Brand, T. M., Iida, M., Li, C. & Wheeler, D. L. The Nuclear Epidermal Growth Factor Receptor Signaling Network and its Role in Cancer. Discov. Med. 12, 419–432 (2011).

20. Tomas, A., Futter, C. E. & Eden, E. R. EGF receptor trafficking: consequences for signaling and cancer. Trends Cell Biol. 24, 26–34 (2014).

21. Chung, B. M. et al. Aberrant trafficking of NSCLC-associated EGFR mutants through the endocytic recycling pathway promotes interaction with Src. BMC Cell Biol. 10, 84 (2009).

22. Huang, W.-C. et al. Nuclear translocation of epidermal growth factor receptor by Akt-dependent phosphorylation enhances breast cancer-resistant protein expression in gefitinib-resistant cells. J. Biol. Chem. 286, 20558–20568 (2011).

23. Ramalingam, S. S. et al. Overall Survival with Osimertinib in Untreated, EGFR-Mutated Advanced NSCLC. N. Engl. J. Med. 382, 41–50 (2020).

24. Kawamata, H., Kameyama, S. & Oyasu, R. In vitro and in vivo acceleration of the neoplastic phenotype of a low-tumorigenicity rat bladder carcinoma cell line by transfected transforming growth factor-alpha. Mol. Carcinog. 9, 210–219 (1994).

25. Ohsaki, Y. et al. Epidermal growth factor receptor expression correlates with poor prognosis in non-small cell lung cancer patients with p53 overexpression. Oncol. Rep. 7, 603–607 (2000).

26. Bethune, G., Bethune, D., Ridgway, N. & Xu, Z. Epidermal growth factor receptor (EGFR) in lung cancer: an overview and update. J. Thorac. Dis. 2, 48–51 (2010).

27. Nicholson, R. I., Gee, J. M. & Harper, M. E. EGFR and cancer prognosis. Eur. J. Cancer Oxf. Engl. 1990 37 Suppl 4, S9–15 (2001).

28. Hou, J. et al. Gene expression-based classification of non-small cell lung carcinomas and survival prediction. PloS One 5, e10312 (2010).

29. Selamat, S. A. et al. Genome-scale analysis of DNA methylation in lung adenocarcinoma and integration with mRNA expression. Genome Res. 22, 1197–1211 (2012).

30. Gao, J. et al. Integrative analysis of complex cancer genomics and clinical profiles using the cBioPortal. Sci. Signal. 6, pl1 (2013).

31. Cerami, E. et al. The cBio cancer genomics portal: an open platform for exploring multidimensional cancer genomics data. Cancer Discov. 2, 401–404 (2012).

32. Cancer Genome Atlas Research Network. Comprehensive molecular profiling of lung adenocarcinoma. Nature 511, 543–550 (2014).

33. Barretina, J. et al. The Cancer Cell Line Encyclopedia enables predictive modelling of anticancer drug sensitivity. Nature 483, 603–607 (2012).

34. Hanahan, D. & Weinberg, R. A. The hallmarks of cancer. Cell 100, 57–70 (2000).

35. Amit, I. et al. A module of negative feedback regulators defines growth factor signaling. Nat. Genet. 39, 503–512 (2007).

36. Mikula, M. et al. Genome-wide co-localization of active EGFR and downstream ERK pathway kinases mirrors mitogen-inducible RNA polymerase 2 genomic occupancy. Nucleic Acids Res. 44, 10150–10164 (2016).

37. Brand, T. M., Iida, M., Li, C. & Wheeler, D. L. The nuclear epidermal growth factor receptor signaling network and its role in cancer. Discov. Med. 12, 419–432 (2011).

38. Yu, Y.-L. et al. Nuclear EGFR suppresses ribonuclease activity of polynucleotide phosphorylase through DNAPK-mediated phosphorylation at serine 776. J. Biol. Chem. 287, 31015–31026 (2012).

