## Supplementary data for "Sortilin exhibits tumor suppressor-like activity by limiting EGFR transducing function"

### Supplementary_Materials_1_Lapeyronnie et al.

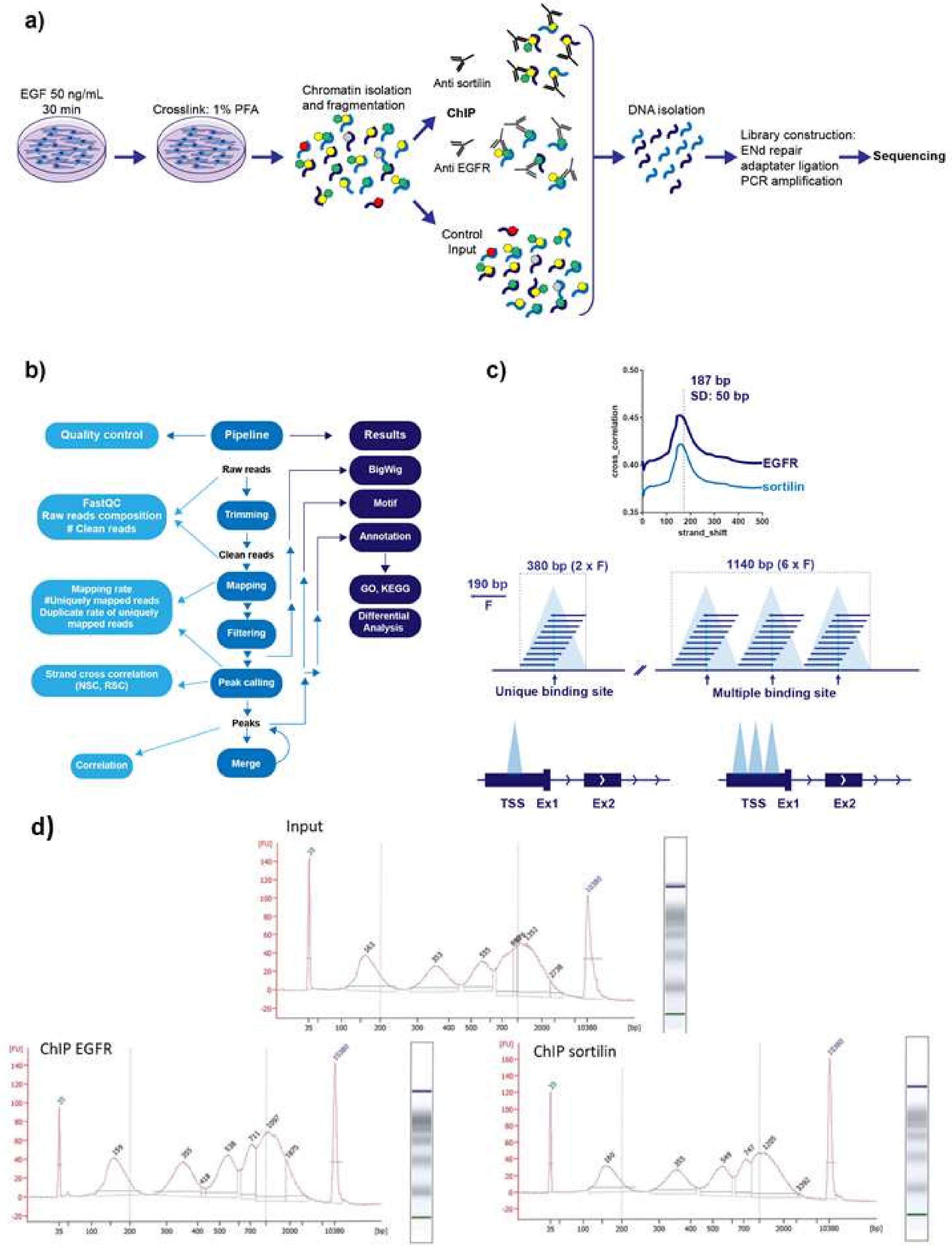

**Supplementary material 1: ChIP workflow and bioinformatic pipeline analysis. (a)** Schematic showing the ChIP protocol from cells to DNA before sequencing. **(b)** Pipeline of bioinformatic analysis. Chromatin immunoprecipitation sequencing (IP-seq) allows identification of the genome regions that interact efficiently with transcription factors and chromatin-associated proteins. Mapping the sequencing results onto the genome can provide genome-wide information about the DNA regions interacting with histones and transcription factors. **(c)** Determination of unique or multiple DNA binding sites on targeted promoter genes. **(d)** Chromatin digestion by microccocal nuclease after ChIP with an Agilent bioanalyzer to verify the presence of mono-nucleosomes in different sample (Input, ChIP EGFR, and ChIP sortilin).

### Supplementary_Figure_1_Lapeyronnie et al.

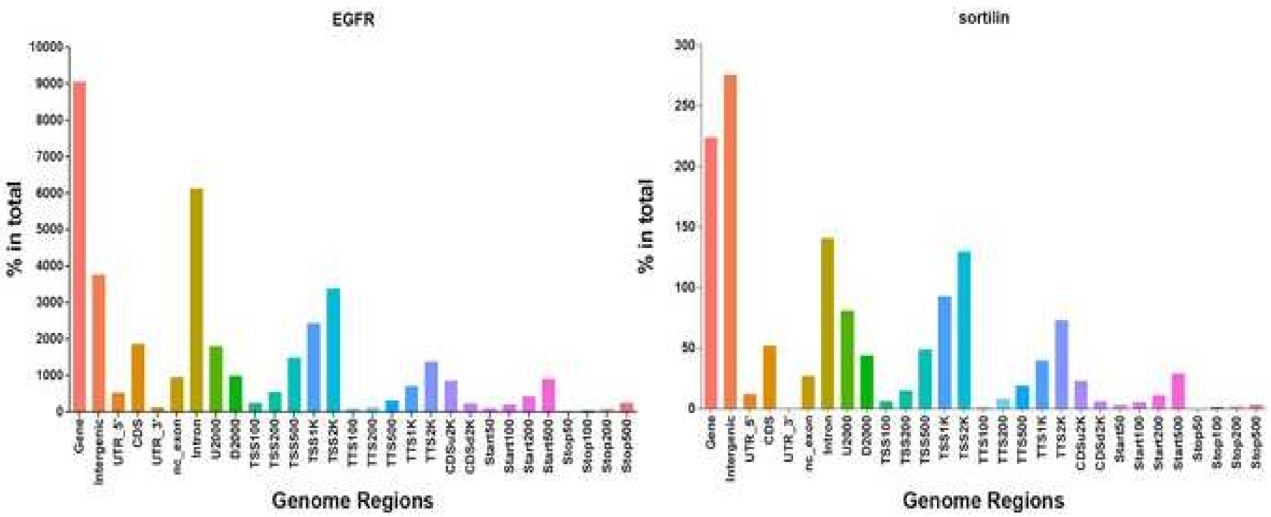


**Supplementary Figure 1: Pathways and genome regions targeted by EGFR and sortilin.** (a-b) Representations of genome regions targeted by (a) EGFR and (b) sortilin. Distribution of peaks in different functional areas. The horizontal axis represents different functional areas, and the vertical axis represents the ratio of the peak in the functional region to the total peaks. The number at the top of each functional region represents the peak number. U2000 and D2000 indicate 2000 bp in the upstream and downstream regions, respectively; CDSu2K and CDSd2K indicate 2 kb upstream and downstream of the CDS, respectively; and TSS100, TTS100, Start100, and Stop100 indicate the 100 bp regions centered on the TSS, TTS, Start-codon, and Stop-codon, respectively.

### Supplementary_Figure_2_Lapeyronnie et al.

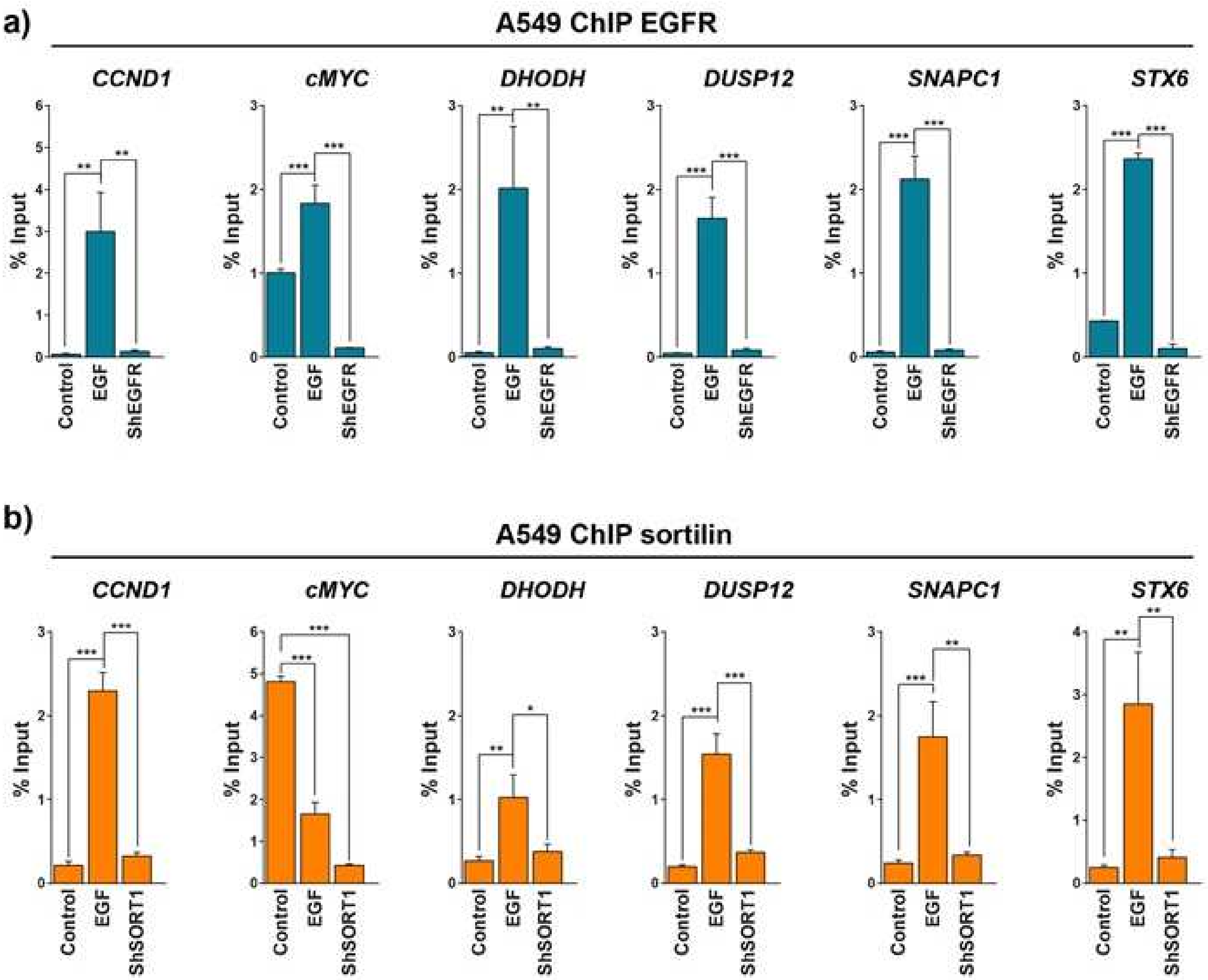


**Supplementary Figure 2: EGF-associated increases in sortilin binding to DNA in A549 cells.** EGFR or sortilin was immunoprecipitated, and qPCR targeting *CCND1*, *cMYC*, *DHODH*, *DUSP12*, *SNPAC1*, and *STX6* sequences was performed. **(a)** ChIP-qPCR assays of A549 cells in the absence of EGF (control) or in the presence of 50 ng/mL EGF (EGF) and of shEGFR cells (ShEGFR) in the presence of 50 ng/mL EGF. qPCR results were normalized relative to non-relevant antibodies and input. **(b)** ChIP-qPCR assays after sortilin ChIP of A549 cells without EGF (control) or in the presence of 50 ng/mL EGF (EGF), and of ShSORT1 cells (ShSORT1) in the presence of 50 ng/mL EGF. All values represent means ± SD; *$p$<0.05, **$p$<0.01, and ***$p$<0.001.

### Supplementary_Figure_3_Lapeyronnie et al.

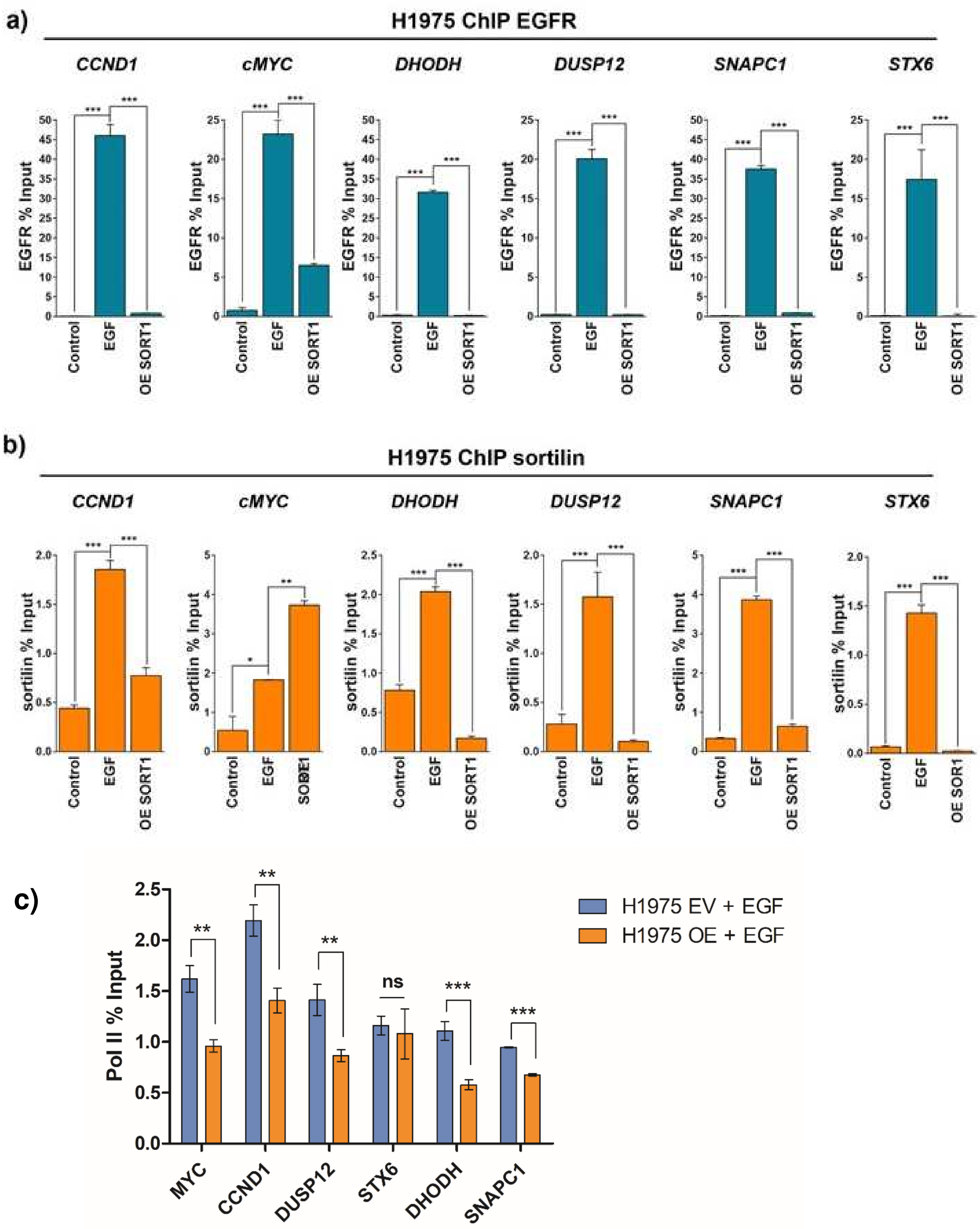


**Supplementary Figure 3: Increase of sortilin binding to DNA in the presence of EGF in H1975 cells and sortilin limits Pol II binding**. qPCR targeting *CCND1*, *cMYC*, *DHODH*, *DUSP12*, *SNPAC1*, and *STX6* sequences was performed after (a) EGFR or (b) sortilin ChIP. (a-b) ChIP-qPCR assays of H1975 cells in the absence (control) or presence of 50 ng/mL EGF (EGF) or *SORT1*

overexpressing cells (OE SORT1) in the presence of 50 ng/mL EGF. qPCR results were normalized relative to non-relevant antibodies and input. **(c)** RNA POLII ChIP assays of H1975 cells expressing empty vector (EV) and overexpressing SORT1 (OE). All values represent means ± SD; *$p<0.05$, **$p<0.01$, and ***$p<0.001$.

#### Supplementary_Figure_4_Lapeyronnie et al.

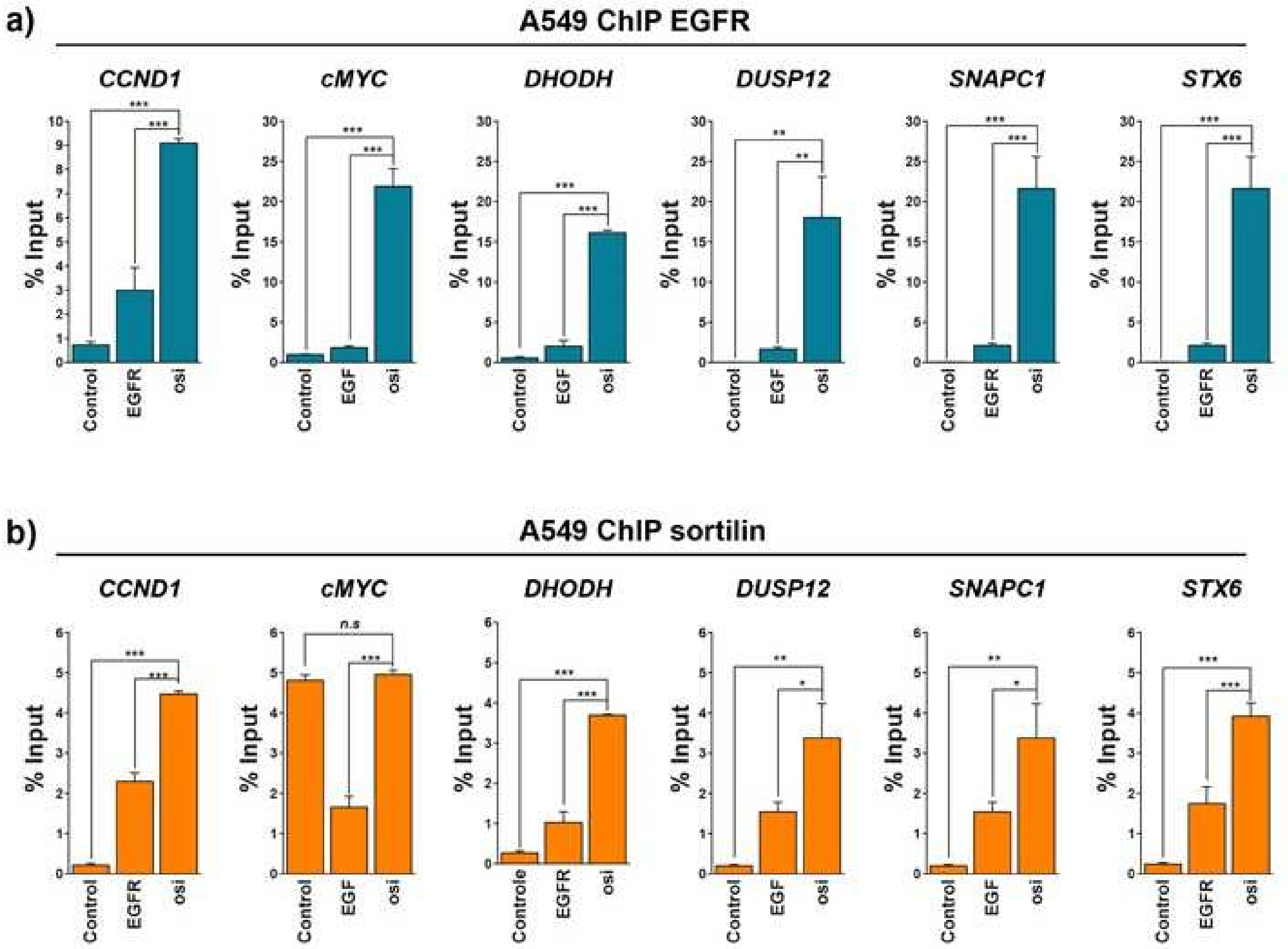


**Supplementary Figure 4: Osimertinib increases EGFR and sortilin binding to DNA in A549 cells.** qPCR amplifications targeting *CCND1*, *cMYC*, *DHODH*, *DUSP12*, *SNPAC1*, and *STX6* sequences performed after (a) EGFR or (b) sortilin ChIP. (a-b) ChIP-qPCR assays of A549 cells in the absence (control) or presence (EGF) of 50 ng/mL EGF or in the presence of 1 µM osimertinib (osi). qPCR results were normalized relative to non-relevant antibodies and input. All values represent means ± SD; **p*<0.05, ***p*<0.01, and ****p*<0.001.

### Supplementary_Figure_5_Lapeyronnie et al.

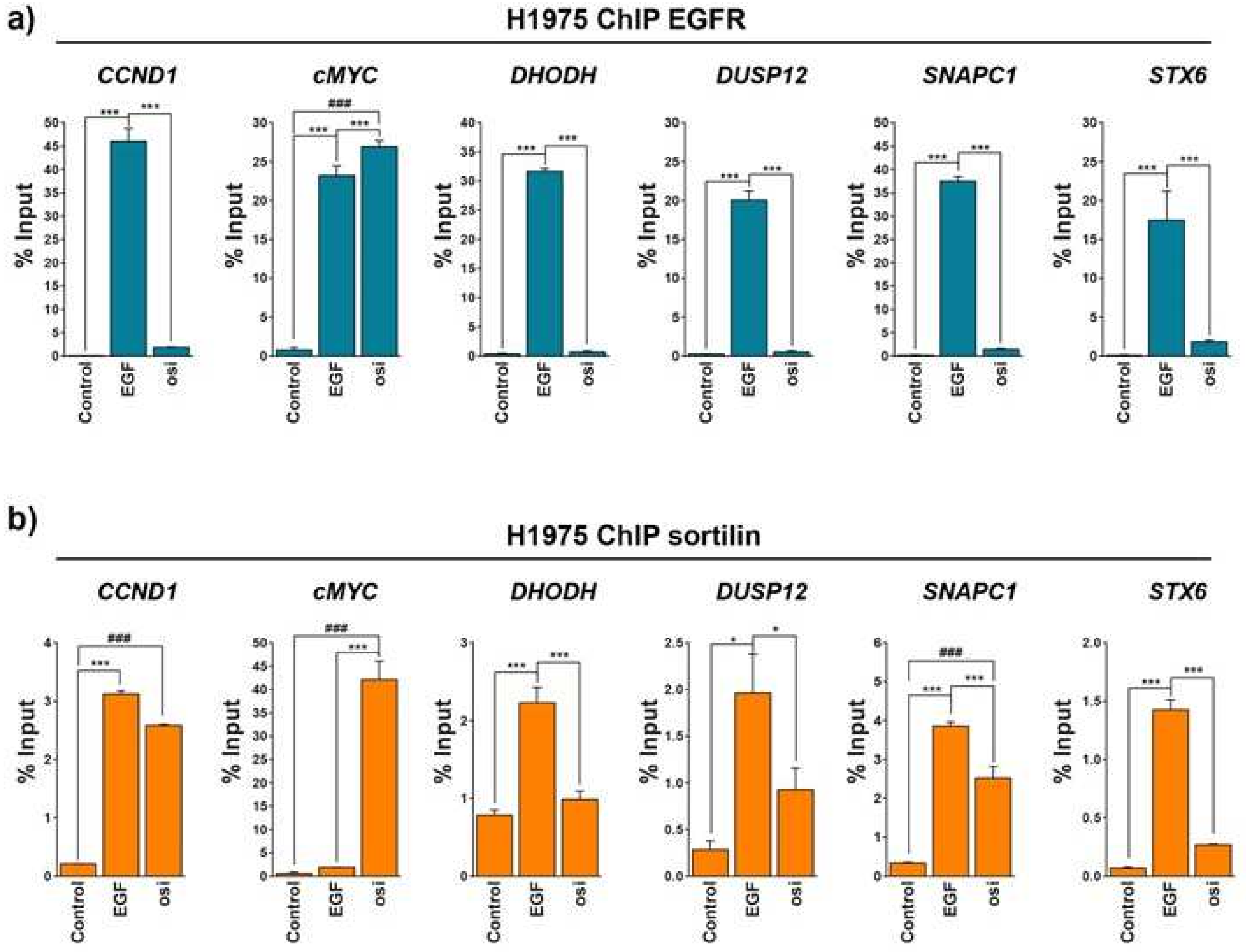


**Supplementary Figure 5: Osimertinib increases EGFR and sortilin binding to DNA in H1975 cells.** qPCRs amplifications targeting *CCND1*, *cMYC*, *DHODH*, *DUSP12*, *SNPAC1*, and *STX6* sequences after (a) EGFR or (b) sortilin ChIP. (a-b) ChIP-qPCR assays of H1975 cells in the absence (control) or presence (EGF) of 50 ng/mL EGF, or in the presence of 1 µM osimertinib (osi). qPCR results were normalized relative to non-relevant antibodies and input. All values represent means ± SD; *$p$<0.05, **$p$<0.01, and ***$p$<0.001.

**Supplementary Table 1.** Gene ontology (GO) analysis of loci shared by EGFR and sortilin pathway components

| GO | Description | Term Type | EGFR *p*-value | Sortilin *p*-value |
|---|---|---|---|---|
| GO:0007156 | Homophilic cell adhesion via plasma membrane adhesion molecule | Biological process | $2.21\ 10^{-11}$ | 0.03 |
| GO:0098742 | Cell–cell adhesion via plasma membrane adhesion molecules | Biological process | $3.67\ 10^{-11}$ | 0.04 |
| GO:0098609 | Cell–cell adhesion | Biological process | $6.70\ 10^{-11}$ | 0.04 |
| GO:0043169 | Cation binding | Molecular function | $2.36\ 10^{-8}$ | 0.03 |
| GO:0046872 | Metal ion binding | Molecular function | $2.87\ 10^{-8}$ | 0.04 |
| GO:0016020 | Membrane | Cellular component | $1\ 10^{-4}$ | 0.04 |
| GO:0038023 | Signaling receptor activity | Molecular function | $4\ 10^{-3}$ | 0.04 |
| GO:0004888 | Transmembrane signaling | Molecular function | $5\ 10^{-3}$ | 0.02 |
| GO:0016021 | Integral component of membrane receptor activity | Cellular component | $9\ 10^{-3}$ | 0.04 |
| GO:0043565 | Sequence-specific DNA binding | Molecular function | 0.017 | 0.01 |
